# NOTCH3 drives meningioma tumorigenesis and resistance to radiotherapy

**DOI:** 10.1101/2023.07.10.548456

**Authors:** Abrar Choudhury, Martha A. Cady, Calixto-Hope G. Lucas, Hinda Najem, Joanna J. Phillips, Brisa Palikuqi, Naomi Zakimi, Tara Joseph, Janeth Ochoa Birrueta, William C. Chen, Nancy Ann Oberheim Bush, Shawn L. Hervey-Jumper, Ophir D. Klein, Christine M. Toedebusch, Craig M. Horbinski, Stephen T. Magill, Aparna Bhaduri, Arie Perry, Peter J. Dickinson, Amy B. Heimberger, Alan Ashworth, Elizabeth E. Crouch, David R. Raleigh

**Author notes:** Authors contributed equally (Abrar Choudhury, Martha A. Cady).

## Abstract

Meningiomas are the most common primary intracranial tumors^1–3^. Treatments for patients with meningiomas are limited to surgery and radiotherapy, and systemic therapies remain ineffective or experimental^4,5^. Resistance to radiotherapy is common in high-grade meningiomas^6^, and the cell types and signaling mechanisms driving meningioma tumorigenesis or resistance to radiotherapy are incompletely understood. Here we report NOTCH3 drives meningioma tumorigenesis and resistance to radiotherapy and find NOTCH3+ meningioma mural cells are conserved across meningiomas from humans, dogs, and mice. NOTCH3+ cells are restricted to the perivascular niche during meningeal development and homeostasis and in low-grade meningiomas but are expressed throughout high-grade meningiomas that are resistant to radiotherapy. Integrating single-cell transcriptomics with lineage tracing and imaging approaches across mouse genetic and xenograft models, we show NOTCH3 drives tumor initiating capacity, cell proliferation, angiogenesis, and resistance to radiotherapy to increase meningioma growth and reduce survival. An antibody stabilizing the extracellular negative regulatory region of NOTCH3^7,8^ blocks meningioma tumorigenesis and sensitizes meningiomas to radiotherapy, reducing tumor growth and improving survival in preclinical models. In summary, our results identify a conserved cell type and signaling mechanism that underlie meningioma tumorigenesis and resistance to radiotherapy, revealing a new therapeutic vulnerability to treat meningiomas that are resistant to standard interventions.

## Main text

Meningiomas arising from the meningothelial lining of the central nervous system comprise more than 40% of primary intracranial tumors^1–3^, and approximately 1% of humans will develop a meningioma in their lifetime^9^. The World Health Organization (WHO) grades meningiomas according to histological features such as mitotic count and rare molecular alterations that are associated with poor outcomes^3^. Surgery and radiotherapy are the mainstays of meningioma treatment^10^, and most WHO grade 1 meningiomas can be effectively treated, but many grade 2 or grade 3 meningiomas are resistant to treatment and cause neurological morbidity and mortality^4^. All clinical trials of systemic therapy have failed to block meningioma growth or improve patient survival^4,5^, and conserved mechanisms underlying aggressive meningiomas remain elusive.

Recent bioinformatic investigations have shed light on meningioma biology, revealing molecular groups of tumors with distinct clinical outcomes that provide a framework for investigating meningioma resistance to treatment^11–18^. Merlin-intact meningiomas encoding the tumor suppressor *NF2* on chromosome 22q are often low-grade, have favorable clinical outcomes, and are sensitive to cytotoxic treatments such as radiotherapy^11^. Meningiomas with biallelic inactivation of *NF2* are often high-grade, have poor clinical outcomes, and can be resistant to radiotherapy^13^. Cancer stem cells can mediate resistance to radiotherapy^19^, but the cell types and signaling mechanisms driving tumorigenesis or resistance to treatment across molecular groups of meningiomas are unknown. Morphologic correlates between meningioma cells and arachnoid cap cells found on meningeal invaginations of dural venous sinuses have fueled a longstanding hypothesis that meningiomas may arise from arachnoid cap cells^20,21^. However, the WHO recognizes 15 histological variants of meningiomas^3^, some of which do not resemble arachnoid cap cells, or which may develop far from dural venous sinuses, including cases of primary intraparenchymal, intraventricular, or pulmonary meningiomas^22–24^. These data suggest additional cell types may contribute to meningioma tumorigenesis, and that understanding cell types and signaling mechanisms underlying meningioma tumorigenesis or resistance to radiotherapy may reveal new targets for systemic therapies to treat patients with meningiomas.

### NOTCH3 is enriched in meningioma mural cells and is expressed throughout high-grade meningiomas

To elucidate the cellular architecture of meningiomas with poor clinical outcomes, single-cell RNA sequencing was performed on 30,934 cells from 6 human meningioma samples with biallelic inactivation of *NF2*, including loss of at least 1 copy of chromosome 22q (Fig. 1a and Extended Data Fig. 1a). Datasets were integrated and corrected for batch effects using Harmony^25^ (Extended Data Fig. 1b), and uniform manifold approximation and projection (UMAP) revealed a total of 13 cell clusters that were defined using automated cell type classification^26^, cell signature gene sets^27^, cell cycle analysis, and differentially expressed cluster marker genes (Supplementary Table 1). Reduced dimensionality clusters of meningioma tumor cells with loss of chromosome 22q were distinguished from microenvironment cells using CONICS^28^ (Fig. 1b and Extended Data Fig. 1c). Meningioma cell clusters expressing markers of cell cycle progression (C0, C3, C4), signal transduction (C1), or extracellular matrix remodeling (C10) grouped together in UMAP space (Fig. 1a, Extended Data Fig. 1d-g, and Supplementary Table 1). Three meningioma cell clusters with loss of chromosome 22q were distinguished from other meningioma cell types by expression of mural cell markers (C7, C11, C12), including genes associated with pericytes (*CD248, ABCC9, CSPG4, GJA4*), fibroblasts (*SERPING1, CLU*), smooth muscle cells (*SDC2, ACTA2*), and multiple mural cell lineages (*PDGRFB, RGS5*)^29–31^ (Fig. 1a-c, Extended Data Fig. 1f, and Supplementary Table 1). The cluster of cells expressing endothelial markers (C8), including genes associated with tip cells (*ADM, ANGPT2, COL9A3*), capillary cells (*MFSD2A*), arterial cells (*CXCL12*), and multiple endothelial cell lineages (*CD34, VWF, CLDN5, PECAM1, PDGFD, KDR, FLT, FLT1, TIE1*)^29–31^, did not show loss of chromosome 22q (Fig. 1a-c, Extended Data Fig. 1f, and Supplementary Table 1).

**Fig. 1.**
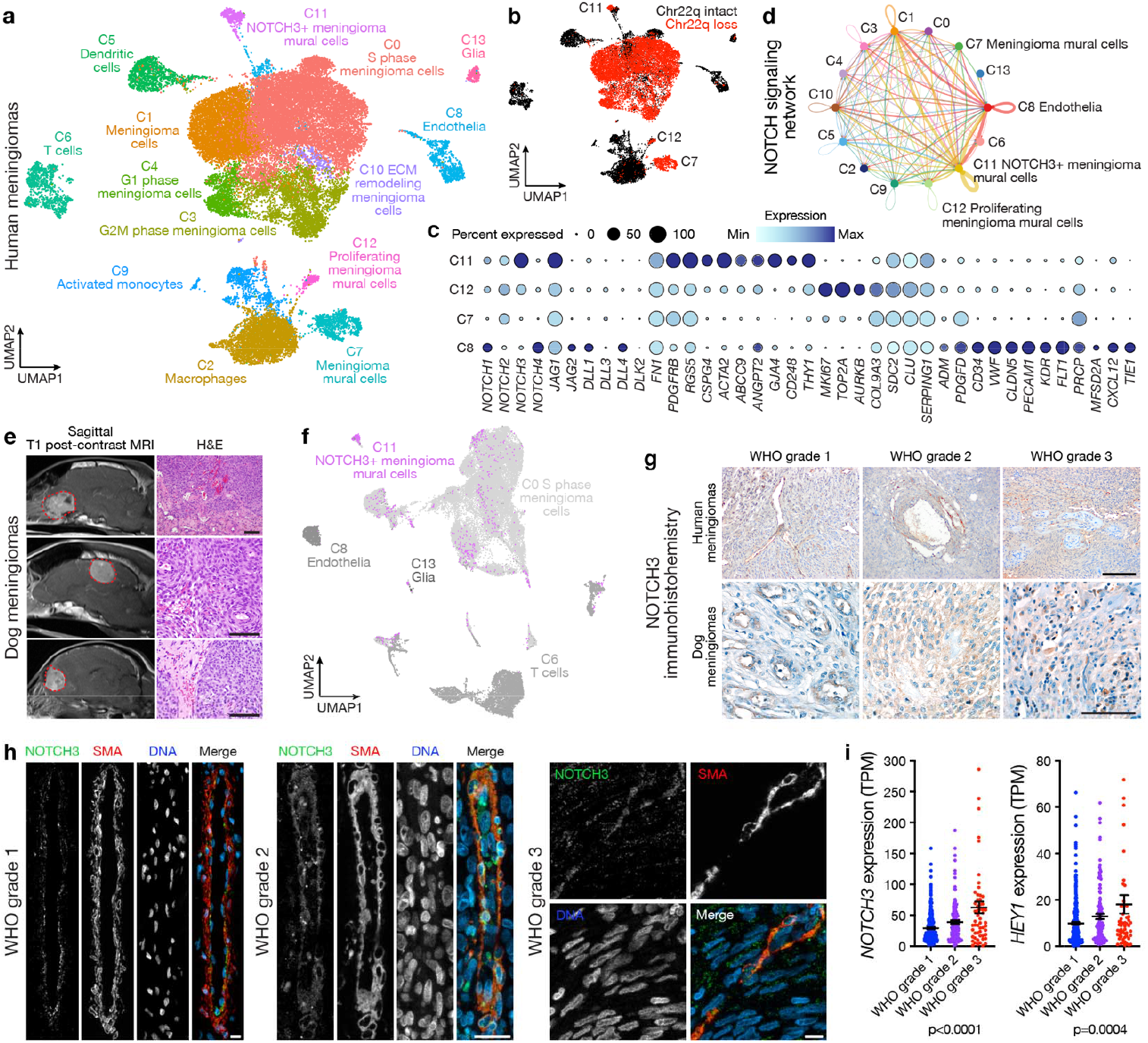
NOTCH3 is enriched in meningioma mural cells and is expressed throughout high-grade meningiomas. **a**, Single-cell RNA sequencing UMAP of 30,934 transcriptomes from human meningioma samples with loss of chromosome 22q showing tumor cell states and microenvironment cell types. **b**, UMAP showing single-cell RNA sequencing of human meningiomas shaded by chromosome 22q status. **c**, Dot plot showing expression of NOTCH receptors (*NOTCH1, NOTCH2, NOTCH3, NOTCH3)*, NOTCH ligands (*JAG1, JAG2, DLL1, DLL3, DLL4, DLK2, FN1)*, mural cell markers (*PDGFRB, RGS5, CSPG4, ACTA2, ABCC9, ANGPT2, GJA4, CD248, COL9A3, SDC2, CLU, SERPING1*), cancer stem-cell marker (*NOTCH3, THY1*), cell proliferation markers (*MKI67, TOP2A, AURKB*), and endothelial cells markers (*ADM, PDGFD, CD3, VWF, CLDN5, PECAM1, KDR, FLT1, PRCP, MFSD2A, CXCL12, TIE1*) across meningioma mural (C7, C11, C12) or endothelial cells (C8) from **a. d**, Inference of NOTCH signaling network in human meningiomas using single-cell RNA sequencing cell-cell communication analysis. **e**, Magnetic resonance imaging (MRI, left) or H&E images (right) of spontaneous dog meningiomas. Scale bars, 100µm. **f**, Transcriptomic concordance of human meningioma single-cell cluster identities from **a** projected on single-cell RNA sequencing UMAP of 40,525 transcriptomes from dog meningioma samples showing NOTCH3+ meningioma mural cells and proliferating meningioma cells are conserved across human and dog meningiomas. **g**, IHC for NOTCH3 across histological grades of human (top) or dog (bottom) meningiomas. Representative of n=3-10 meningiomas per grade. Scale bars, 100µm. **h**, IF for NOTCH3 and the mural cell marker SMA across histological grades of human meningiomas. DAPI marks DNA. Representative of n=10 meningiomas per grade. Scale bars, 10µm. **i**, Quantification of *NOTCH3* or the NOTCH3 target gene *HEY1* across meningioma grades using RNA sequencing of n=502 human meningiomas. TPM, transcripts per million. Lines represent means and error bars represent standard error of means. ANOVA.

Meningioma mural cell clusters were distinguished from one another by expression of markers of cell cycle progression (*MKI67, TOP2A, AURKB*) or markers of cancer stem cells (*NOTCH3, THY1*)^32–34^ (Fig. 1c). The cluster of meningioma mural cells that was enriched in *NOTCH3* and *THY1* (C11), and the endothelial cell cluster (C8), also expressed NOTCH ligands (*JAG1, JAG2, DLL1, DLL4, FN1*) (Fig. 1c). Cell-cell communication analysis of single-cell RNA sequencing data using CellChat^35^ suggested the NOTCH signaling network was active in meningiomas with loss of chromosome 22q, particularly among and between NOTCH3+ meningioma mural cells and endothelia (Fig. 1d). Moreover, *NOTCH3* was expressed in meningioma cell clusters marked by the meningioma gene *SSTR2A*^36^ (Extended Data Fig. 1h) but other NOTCH receptors were not enriched in meningioma cell types (Supplementary Table 1).

The evolutionarily conserved NOTCH family of transmembrane proteins enable intercellular communication to regulate mammalian cell fate and growth^37–39^. Similar to humans, the most common primary intracranial tumors in dogs are meningiomas^40^. Thus, to determine if NOTCH3+ meningioma mural cells were conserved in meningiomas from other mammals, single-cell RNA sequencing was performed on 40,525 cells from 5 samples from dog meningiomas (Fig. 1e and Extended Data Fig. 2a). Datasets were integrated and corrected for batch effects using Harmony^25^ (Extended Data Fig. 2b), and UMAP revealed a total of 14 cell clusters that were defined using automated cell type classification^26^, cell signature gene sets^27^, cell cycle analysis (Extended Data Fig. 2c), and differentially expressed cluster marker genes (Extended Data Fig. 2d-f and Supplementary Table 2). Human meningioma single-cell cluster identities were projected onto reduced dimensionality clusters of dog meningioma cells using transcriptional correlation and expression of conserved marker genes across species, revealing conservation of endothelia, glia, and immune cell types (Fig. 1f). NOTCH3+ meningioma mural cells from human meningiomas were found in a unique dog meningioma cluster of mural cells and a mixed cluster that was also comprised of cycling meningioma cells from human meningiomas (Fig. 1f). These data suggest NOTCH3+ meningioma mural cells are conserved in mammalian meningiomas. In support of this hypothesis, immunohistochemistry (IHC) for NOTCH3 across all grades of human or dog meningiomas showed NOTCH3 was heterogeneous and predominantly restricted to the perivascular niche in low-grade meningiomas but was expressed throughout high-grade meningiomas from both species (Fig. 1g and Extended Data Fig. 3). Immunofluorescence (IF) across all grades of human meningiomas demonstrated that NOTCH3 was expressed by mural cells marked by SMA and was enriched throughout high-grade meningiomas but was not expressed by endothelial cells marked by VWF (Fig. 1h and Extended Data Fig. 4-9). RNA sequencing of 502 human meningiomas^11,13^ showed that expression of *NOTCH3* and the NOTCH3 target gene *HEY1* increased from WHO grade 1 (n=329) to grade 2 (n=117) to grade 3 (n=56) meningiomas.

### NOTCH3+ mural cells underlie meningeal tumorigenesis

*NOTCH3* mutations in humans cause CADASIL^41^, a hereditary adult-onset cerebral arteriopathy associated with sub-cortical ischemic events and alterations in brain vascular smooth muscle cells^42^. The functional relevance of NOTCH3 for meningeal development, homeostasis, or tumorigenesis is unknown. IF of the developing human cortex from gestational week 17 showed NOTCH3 was expressed by mural cells marked by PDGFRβ in the cortical plate and margin zone (Extended Data Fig. 10a), both of which contribute to meningeal development^43–46^. Re-analysis of single-cell RNA sequencing data from perinatal human brain vasculature (139,134 cells from gestational weeks 15, 17, 18, 20, 22, and 23)^29^ or adult human brain vasculature (84,138 cells or 52,023 cells from 2 studies)^30,31^ demonstrated *NOTCH3* was enriched in mural cells marked by *PDGFRβ* or *ACTA2* (SMA) (Extended Data Fig. 10b-d). In contrast to *NOTCH3* expression in brain vasculature and meningioma mural cells (Fig. 1c), IHC and IF of adult human meninges showed NOTCH3 expression was restricted to mural cells that were adjacent abut non-overlapping with vascular smooth muscle cells expressing SMA (Fig. 2a, b). Thus, NOTCH3+ mural cells in the brain (Extended Data Fig. 10a-d), meninges (Fig. 2a, b), and meningiomas (Fig. 1 and Extended Data 1-9) have partially overlapping gene and protein expression programs, suggesting these cell types may fulfill different functions during development, homeostasis, or tumorigenesis.

**Fig. 2.**
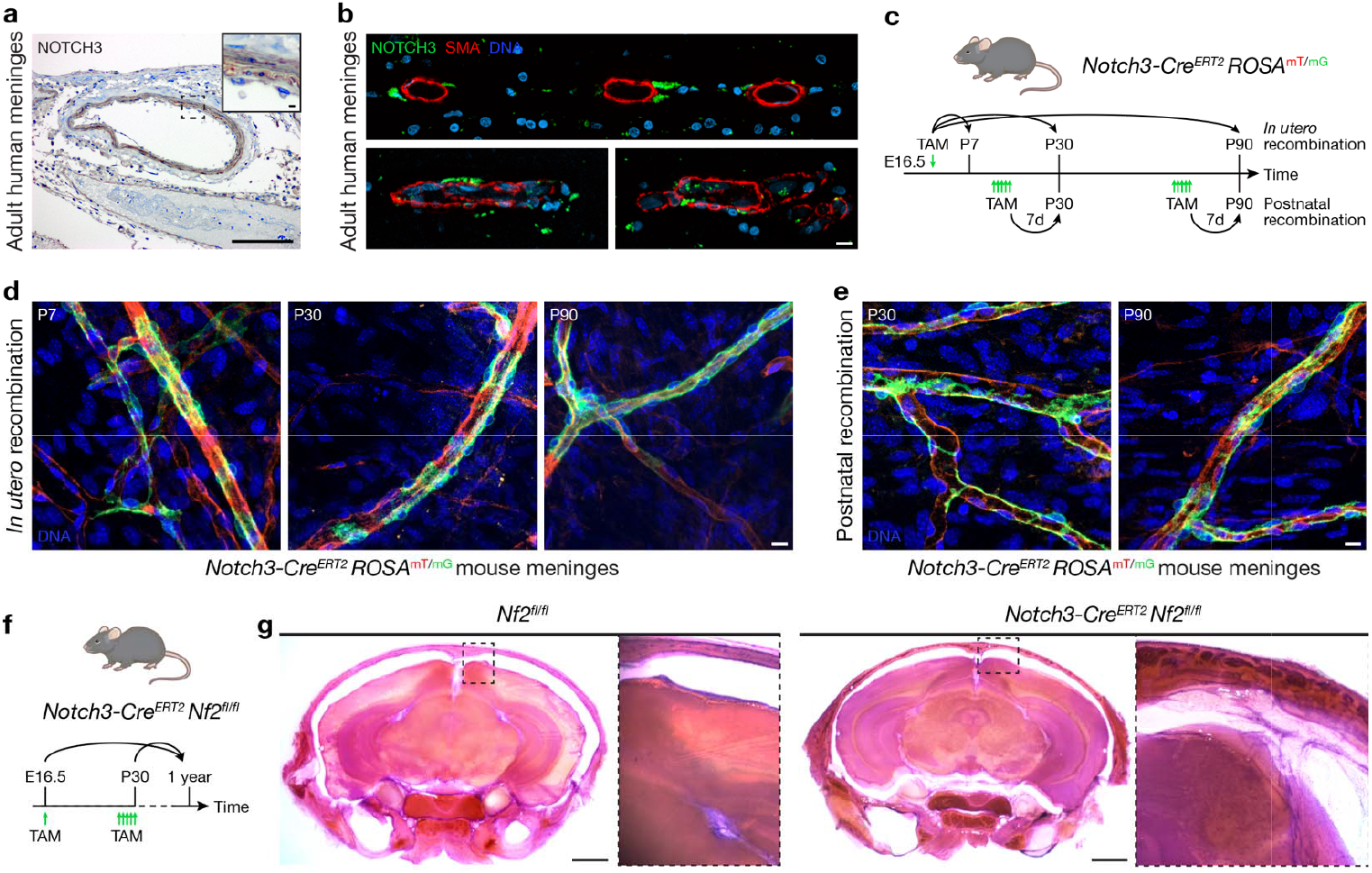
NOTCH3+ mural cells underly meningeal tumorigenesis. **a**, IHC for NOTCH3 in the adult human meninges showing expression is restricted to mural cells. Representative of n=3 biological replicates. Scale bars, 100µm and 10µm (insert). **b**, IF for NOTCH3 and the mural cell marker SMA in 3 adult human meningeal samples showing NOTCH3 is expressed in mural cells adjacent to smooth muscle cells in the meninges. DAPI marks DNA. Scale bar, 10µm. **c**, Experimental design for *in vivo* lineage tracing of NOTCH3+ mural cells during meningeal development (*in utero* recombination) or homeostasis (postnatal recombination). TAM, tamoxifen. **d**, Confocal microscopy of whole mount mouse convexity meningeal samples at P7, P30, or P90 after *in utero* recombination of the *ROSA*^mT/mG^ allele showing NOTCH3 cells (green) are restricted to the perivascular niche during meningeal development. Representative of n=3 biological replicates per timepoint. DAPI marks DNA. Scale bar, 10µm. **e**, Confocal microscopy of whole mount mouse convexity meningeal samples at P30 or P90 after postnatal recombination of the *ROSA*^mT/mG^ allele showing NOTCH3+ cells (green) are restricted to the perivascular niche during meningeal homeostasis. Representative of n=3 biological replicates per timepoint. DAPI marks DNA. Scale bar, 10µm. **f**, Experimental design for *in vivo* biallelic inactivation of *Nf2* in NOTCH3+ cells during meningeal development (E16.5) or homeostasis (P30). Mice were monitored for 1 year after *Nf2* inactivation. **g**, Coronal H&E images of 300µm decalcified mouse skull sections 1 year after treatment of mice with TAM. No gross tumors were identified, but insets show *Nf2* inactivation in NOTCH3+ cells is associated with meningeal hyperproliferation after either *in utero* (E16.5) or postnatal (P30) treatment with TAM. Representative of n=5-8 biological replicates per condition. Scale bars 1mm.

To determine if NOTCH3+ cells give rise to other cell types during meningeal development or homeostasis, a tamoxifen-inducible *Notch3-Cre*^*ERT2*^ allele^47^ was combined with the global double-fluorescent *ROSA*^mT/mG^ Cre reporter allele^48^ (Fig. 2c). Fluorophore recombination was induced during meningeal development (E16.5) or homeostasis (P30, P90) and confocal microscopy of whole mount mouse meningeal samples revealed NOTCH3+ cells were restricted to the perivascular niche across all contexts (Fig. 2d, e). To determine if NOTCH3+ cells underlie meningeal tumorigenesis, the *Notch3-Cre*^*ERT2*^ allele was combined with *Nf2*^fl/fl^ alleles for conditional biallelic inactivation of the *Nf2* tumor suppressor in mice^49^. *Nf2* was inactivated in NOTCH3+ cells *in utero* (E16.5) or in adulthood (P30), and mice were monitored without evidence of neurological symptoms or other phenotypes for 1 year (Fig. 2f). Morphologic examination of the central nervous system revealed meningeal hyperproliferation after biallelic inactivation of *Nf2* in NOTCH3+ cells compared to mice with intact *Nf2* (Fig. 2g). These data support the hypothesis that NOTCH3+ mural cells underlie meningeal tumorigenesis but do not contribute to meningeal self-renewal (Fig. 2d, e), suggesting that beyond being a marker of meningioma initiation, NOTCH3 may represent a therapeutic vulnerability to treat meningiomas that are resistant to standard interventions.

### NOTCH3 signaling drives meningioma tumor initiating capacity, cell proliferation, and angiogenesis

To identify reagents for mechanistic and functional interrogation of NOTCH3 signaling in the context of meningioma tumorigenesis, reference transcriptomic signatures of human meningioma single-cell clusters (Fig. 1a) were used to estimate the proportion of NOTCH3 meningioma mural cells across 502 human meningiomas with matched RNA sequencing and DNA methylation profiling^11,13^. DNA methylation profiling reveals meningiomas are comprised of Merlin-intact, Immune-enriched, and Hypermitotic molecular groups, and can identify meningioma cell lines that are representative of each group^11^. Immune-enriched meningiomas are distinguished from other molecular groups by hypomethylation of genes involved in vasculature development^11^. Cell type deconvolution across meningioma DNA methylation groups showed Immune-enriched meningiomas (n=180) were enriched in NOTCH3 meningioma mural cells compared to Merlin-intact (n=176) or Hypermitotic meningiomas (n=146) (Fig. 3a) and immunoblots demonstrated NOTCH3 was expressed in CH-157MN^50^ and IOMM-Lee^51^ Immune-enriched meningioma cell lines (Fig. 3b).

**Fig. 3.**
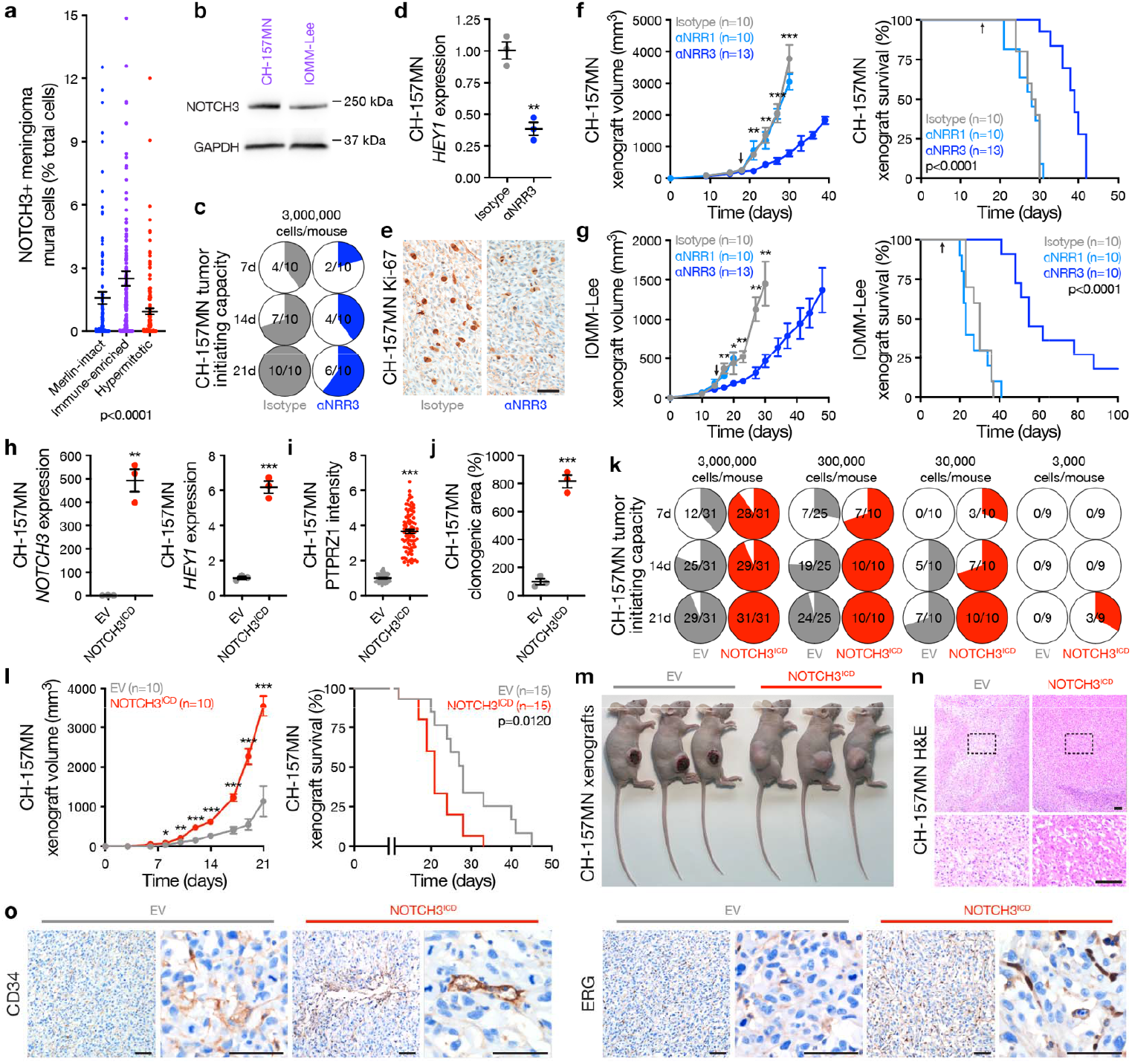
NOTCH3 signaling drives meningioma tumor initiating capacity, cell proliferation, and angiogenesis. **a**, Deconvolution of NOTCH3+ meningioma mural cells from Fig. 1a using human meningiomas with paired RNA sequencing and DNA methylation profiling (n=502). ANOVA. **b**, Immunoblots showing NOTCH3 is expressed in CH-157MN and IOMM-Lee Immune-enriched meningioma cell lines. **c**, *In vivo* tumor initiating capacity of CH-157MN meningioma cells in NU/NU mice ± αNRR3 IP injection 2 times per week. Denominators indicate number of mice at each time point. Numerators indicate number of mice with tumors at each time point. **d**, QPCR for the NOTCH3 target gene *HEY1* from meningioma xenografts ± αNRR3 treatment for 2 weeks. Student’s t test. **e**, IHC for Ki-67 in meningioma xenografts showing αNRR3 blocks meningioma cell proliferation. Representative of n=3 xenografts per condition. Scale bar, 100µm. **f**, CH-157MN meningioma xenograft growth (left, student’s t tests) or survival (log-rank test). Arrows indicate initiation of bi-weekly treatment with the indicated therapy, which continued until death. **g**, IOMM-Lee meningioma xenograft growth (left, student’s t tests) or survival (log-rank test). Arrows as in **f. h**, QPCR for *NOTCH3* or *HEY1* in CH-157MN meningioma cells ± stable expression of empty vector (EV) or NOTCH3^ICD^. Student’s t tests. **i**, IF quantification of the stem cell marker PTPRZ1 in CH-157MN meningioma cells. Student’s t test. **j**, Clonogenic *in vitro* growth of CH-157MN meningioma cells after 2 weeks. Student’s t test. **k**, *In vivo* tumor initiating capacity of CH-157MN meningioma cells ± EV or NOTCH3^ICD^ over limiting dilutions. Numerator and denominator as in **c. l**, CH-157MN meningioma xenograft growth (left, student’s t tests) or survival (log-rank test). **m**, Images of heterotopic meningioma xenografts showing macroscopic necrosis and ulceration in EV meningiomas. Representative of n=7-9 xenografts per condition. **n**, H&E low and high (box) magnification images of meningioma xenografts showing microscopic necrosis in EV meningiomas. Representative of n=3 xenografts per condition. Scale bars, 100µm. **o**, IHC for endothelia markers in meningioma xenografts showing NOTCH3^ICD^ induces meningioma angiogenesis. Representative of n=3 xenografts per condition. Scale bars, 100µm. Lines represent means and error bars represent standard error of means. **p≤0.01, ***p≤0.0001.

NOTCH receptor activation requires ADAM protease cleavage of the extracellular negative regulatory region (NRR), intramembrane proteolysis by the γ-secretase complex, and release of the NOTCH intracellular domain (ICD) that regulates mammalian cell fate and growth^37–39^. Small molecule inhibitors of ADAM or γ-secretase do not distinguish between individual NOTCH receptors and have not been adopted in routine clinical practice due to toxicity, but antibody stabilization of the NRR allows for selective inhibition of individual NOTCH receptors in preclinical models^7^. An antibody selectively stabilizing the NRR of NOTCH3 (αNRR3)^8^ attenuated *in vivo* tumor initiating capacity (Fig. 3c), blocked expression of *HEY1* (Fig. 3d), and reduced cell proliferation of CH-157MN xenografts (Fig. 3e). Moreover, αNRR3 blocked tumor growth and improved survival of CH-157MN (Fig. 3f) and IOMM-Lee xenografts (Fig. 3g) compared to isotype control treatment or treatment with αNRR1. Overexpression of the NOTCH3 ICD (NOTCH3^ICD^) increased expression of *HEY1* and PTPRZ1, a marker of meningioma self-renewal^52^, and increased clonogenic growth of CH-157MN cells *in vitro* compared to CH-175MN cells expressing empty vector (EV) control (Fig. 3h-j). NOTCH3^ICD^ increased the tumor initiating capacity of CH-157MN cells using *in vivo* limiting dilution assays (Fig. 3k), and increased tumor growth and reduced survival of CH-157MN xenografts compared to EV (Fig. 3l). Meningiomas are not protected by the blood brain barrier^53^, and heterotopic CH-157MN EV xenografts developed ulceration and necrosis that was not detected with overexpression of NOTCH3^ICD^ (Fig. 3m, n), suggesting NOTCH3 may contribute to meningioma angiogenesis. In support of this hypothesis, immunostaining for endothelial cell markers (CD34, ERG) was increased in CH-157MN meningioma xenografts with overexpression of NOTCH3^ICD^ compared to EV (Fig. 3o).

### NOTCH3 signaling drives meningioma resistance to radiotherapy

To determine if NOTCH3 underlies meningioma recurrence after standard interventions, differential expression (Supplementary Table 3) and gene ontology analyses (Fig. 4a) were performed on RNA sequencing data from primary (n=403) compared to recurrent human meningiomas after treatment with surgery and radiotherapy (n=99)^11,13^. Recurrent meningiomas were distinguished by gene expression programs controlling DNA metabolism, radiotherapy response, cell signaling, and cell proliferation (Fig. 4a and Supplementary Table 3). Immunostaining for the cell proliferation marker Ki-67, deconvolved NOTCH3 meningioma mural cells from RNA sequencing, and expression of *NOTCH3* and *HEY1* were enriched in recurrent compared to primary meningiomas (Fig. 4b). Multiplexed sequential IF (seqIF) on 4 pairs of patient-matched meningiomas that were treated with radiotherapy between initial and salvage resections showed NOTCH3 and Ki-67 were enriched in recurrent compared to primary tumors (Fig. 4c and Supplementary Table 4). In preclinical models, radiotherapy attenuated the growth of CH-157MN EV xenografts but did not attenuate the growth of CH-157MN NOTCH3^ICD^ xenografts, which had worse survival than CH-157MN EV xenografts despite treatment of both models with ionizing radiation (Fig. 4d).

**Fig. 4.**
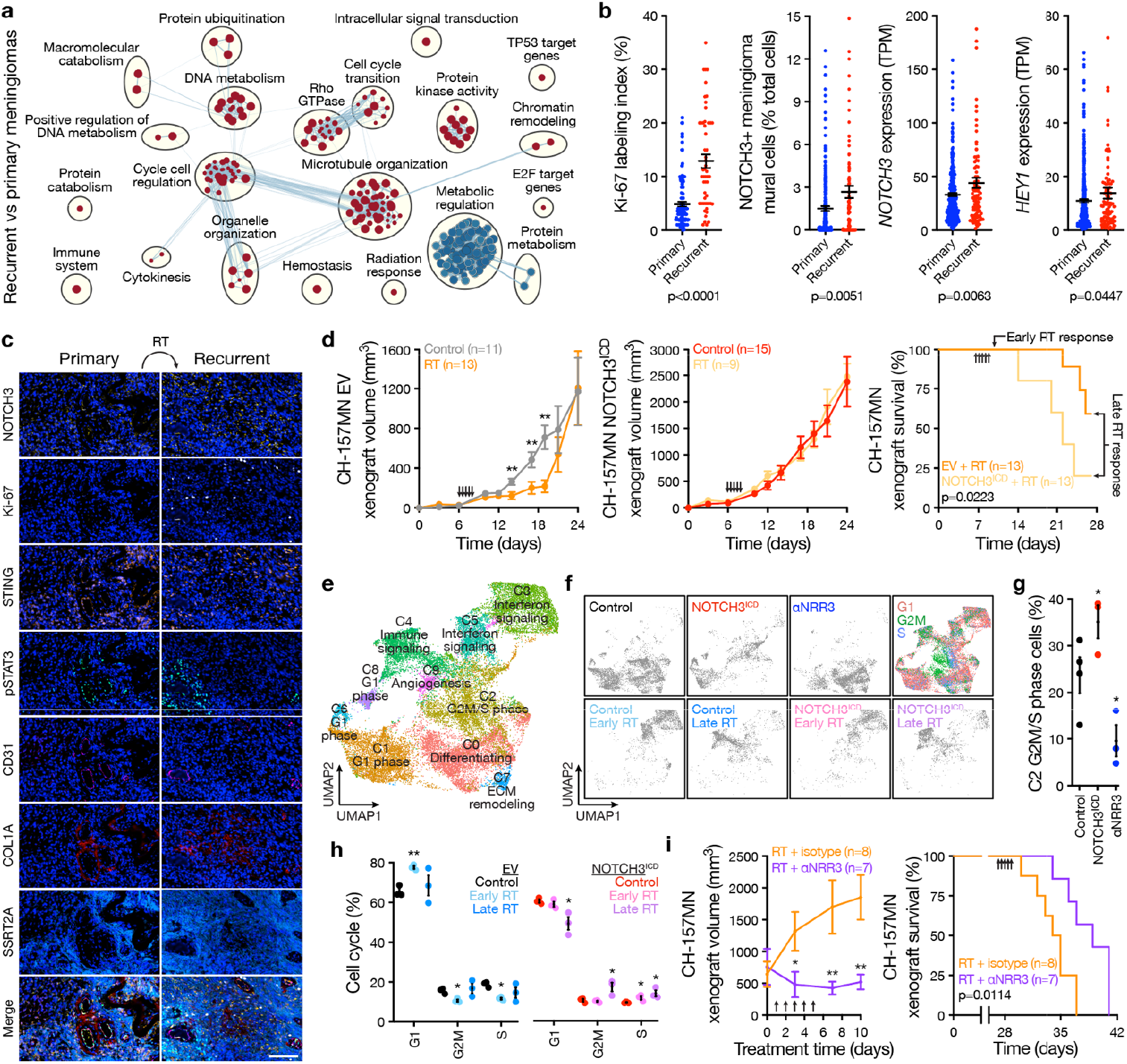
NOTCH3 signaling drives meningioma resistance to radiotherapy. **a**, Network of gene circuits distinguishing recurrent (n=99) from primary (n=403) human meningiomas using RNA sequencing. Nodes represent pathways and edges represent shared genes between pathways (p≤0.01, FDR≤0.01). Red nodes are enriched and blue nodes are suppressed in recurrent versus primary meningiomas. **b**, IHC for Ki-67 in recurrent (n=53) versus primary (n=123) meningiomas, or RNA sequencing of recurrent (n=99) versus primary (n=403) meningiomas for deconvolution of NOTCH3+ meningioma mural cells from Fig. 1a or quantification of *NOTCH3* or *HEY1* expression. TPM, transcripts per million. ANOVA. **c**, Multiplexed seqIF microscopy showing human meningioma recurrence after radiotherapy (RT) is associated with increased NOTCH3 and Ki-67. Many NOTCH3+ cells also express the interferon and innate immune regulators STING and pSTAT3. CD31 marks pericytes, COL1A marks fibroblasts, SSTR2A marks meningioma cells, and DAPI marks DNA. Representative of n=4 pairs of patient-matched primary and recurrent meningiomas. Scale bar, 100µm. **d**, CH-157MN meningioma xenograft growth (left and middle, student’s t tests) or survival (log-rank test) after expression of empty vector (EV) or NOTCH3^ICD^ ± RT showing NOTCH3 drives resistance to RT. Arrows indicate RT treatments (2Gy x 5 daily fractions). Xenografts from all arms were isolated for single-cell RNA-sequencing 1 day after completing RT (early) or once median survival was reached in the EV + RT arm (late). **e**, Single-cell RNA sequencing UMAP of 152,464 meningioma xenograft human cell transcriptomes showing tumor cell states ± αNRR3 treatment for 2 weeks as in Fig. 3f or ± NOTCH3^ICD^ ± RT as in **d. f**, UMAP showing single-cell RNA sequencing of meningioma xenograft human cells shaded by experimental condition or phase of the cell cycle. **g**, Analysis of C2 G2M/S phase meningioma xenograft human cells in control versus NOTCH3^ICD^ versus αNRR3 conditions showing NOTCH3 drives meningioma cell proliferation. Colors as in **f**. Student’s t tests. **h**, Cell cycle analysis across all clusters of meningioma xenograft human cells ± NOTCH3^ICD^ ± RT showing NOTCH3 sustains cell proliferation through G2M and S phase despite RT. Student’s t test. **i**, Meningioma xenograft growth (left, student’s t tests) or survival (log-rank test) after treatment with RT as in **d** ± αNRR3 as in Fig. 3f. αNRR3 treatment was initiated on the first day of radiotherapy and continued until death. Lines represent means and error bars represent standard error of means. *p<0.05, **p≤0.01.

Single-cell RNA sequencing was performed on CH-157MN xenografts after αNRR3 compared to isotype control treatment (Fig. 3f), or after radiotherapy compared to control treatment with EV or NOTCH3^ICD^ overexpression (Fig. 4d). Xenografts were isolated for single-cell RNA sequencing the day after completing radiotherapy (i.e. early) or once median survival was reached in the control arm (i.e. late) to interrogate early versus late effects of ionizing radiation on meningioma cell types (Fig. 4d). Single-cell transcriptomes were mapped to the human and mouse genomes, revealing 152,464 human meningioma cells and 35,230 mouse microenvironment cells across 8 conditions and 23 biological replicates (Extended Data Fig. 11a-e). Reduced dimensionality cell clusters were defined using automated cell type classification^26^, cell signature gene sets^27^, cell cycle analysis (Extended Data Fig. 11b), and differentially expressed cluster marker genes (Extended Data Fig. 12a-e and Supplementary Table 5, 6). Genetic activation (NOTCH3^ICD^) or pharmacologic inhibition (αNRR3) of NOTCH3 influenced the neutrophil, fibroblast, Langerhans cell, conventional dendritic cell, NK cell, and mural cell composition of meningioma xenografts (Extended Data Fig. 12c-e), and NOTCH3+ cells in human meningiomas analyzed using multiplexed seqIF expressed regulators of interferon signaling and innate immune responses (STING, pSTAT3) (Fig. 4c). Radiotherapy induced early monocyte infiltration (Extended Data Fig. 12c-e) and increased interferon and innate immune gene expression from meningioma cells that diminished over time but was conserved across NOTCH3^ICD^ and EV conditions (Fig. 4e, f and Extended Data Fig. 12a, b). Cell cycle analysis across single-cell clusters revealed meningioma cell proliferation was increased by NOTCH3^ICD^ and inhibited by αNRR3 (Fig. 4g). Moreover, radiotherapy inhibited meningioma cell cycle progression in EV xenografts, but NOTCH3^ICD^ sustained the cell cycle through G2M and S phase and increased cell cycle progression in meningioma xenograft samples from late timepoints (Fig. 4h). These data suggest NOTCH3 drives meningioma resistance to radiotherapy through cell cycle progression. In support of this hypothesis, radiotherapy in combination with αNRR3 was more effective than radiotherapy alone at blocking the growth and improving survival from CH-157MN xenografts (Fig. 4i).

## Discussion

Here we report NOTCH3 drives meningioma tumorigenesis and resistance to radiotherapy in cell types that are conserved across meningiomas from humans, dogs, and mice. Our results reveal a new therapeutic vulnerability to treat meningiomas that are resistant to standard interventions, and more broadly suggest that meningioma mural cells may be an effective target to treat the most common primary intracranial tumor.

Our data suggest meningioma vasculature is comprised of endothelia from the microenvironment and tumor cells that may fulfill mural cell functions (Fig. 1a-c). NOTCH3 signaling between meningioma mural cells and endothelia (Fig. 1d) and NOTCH3 mediated intratumor angiogenesis (Fig. 3o) may contribute to meningioma migration into surrounding tissues. In support of this hypothesis, meningiomas without evidence of direct brain parenchyma invasion can nevertheless migrate into Virchow-Robin perivascular cavities that surround perforating arteries and veins in the brain (Extended Data Fig. 13a-c). This unique pattern of migration suggests microscopic positive margins along brain vasculature may contribute to meningioma recurrence after standard interventions.

NOTCH3+ meningioma mural cells demonstrate several hallmarks of cancer stem cells^19^, such as driving meningeal (Fig. 2g) or meningioma cell proliferation (Fig. 3e, 4g, 4h), clonogenic growth (Fig. 3j) or tumor initiating capacity (Fig. 3c, k), angiogenesis (Fig. 3m-o), and resistance to treatment (Fig. 4d, i). NOTCH3 marks cancer stem cells in lung, colon, and breast cancers^32,54–56^, and despite broad NOTCH3 expression in meningiomas, only NOTCH3 meningioma mural cells express other cancer stem cell markers, such as *THY1*^33,34^ (Extended Data Fig. 1h). Our study focuses on meningiomas with loss of *NF2*, and considering the histological^3^ and anatomical diversity of meningiomas^22–24^, it is likely that other stem or progenitor cells contribute to meningioma tumorigenesis, particularly for Merlin-intact meningiomas. PTGDS has been proposed as a marker of meningioma progenitor cells^57^, and while *PTGDS* was only expressed in a subset of single-cells from our study (Extended Data Fig. 1h), *PTGDS* is more broadly expressed throughout Merlin-intact meningioma single-cell clusters^58^. Fluorescent lineage tracing using a *Ptgds-Cre* allele^57^ shows PTGDS cells are not restricted to the perivascular stem cell nice in the meninges of mice (Extended Data Fig. 14). Thus, if meningiomas from different molecular or histological groups or from different anatomic locations can arise from different stem or progenitor cells, it is possible that the vascular phenotypes we report may be unique to NOTCH3 meningioma mural cells.

*NOTCH3* and NOTCH3 target genes are enriched in high-grade (Fig. 1i) and recurrent meningiomas (Fig. 4b), and NOTCH3+ meningioma mural cells are enriched in meningiomas from the Immune-enriched meningioma DNA methylation group (Fig. 3a), but we identify NOTCH3 signaling and NOTCH3+ cells across all grades and molecular groups of meningiomas. Meningiomas from the Hypermitotic DNA methylation groups have the worst clinical outcomes, highest rate of recurrence, and are distinguished by convergent genetic and epigenetic mechanisms that misactivate the cell cycle^11^. RNAScope shows enriched and diffuse *NOTCH3* expression in recurrent Hypermitotic meningiomas (Extended Data Fig. 15). Thus, it is possible that meningiomas from multiple WHO grades or DNA methylation groups with poor clinical outcomes may benefit from treatment with αNRR3 depending on clinical presentation and prior therapy.

The safety and efficacy of selective NOTCH3 inhibition has not been defined in humans, but *Notch3* knockout mice are viable and fertile^59^, and we found that diphtheria toxin ablation of NOTCH3+ cells *in utero* or in adulthood using the *Notch3-Cre*^*ERT2*^ and *ROSA*^iDTR^ alleles^60^ was not associated with neurological symptoms or other phenotypes. In contrast to αNRR3, αNRR1 did not block the growth of meningioma xenografts (Fig. 3f, g) and was associated with skin rash and diarrhea leading to weight loss (Extended Data Fig. 16a). A small molecule inhibitor of the γ-secretase complex blocked meningioma xenograft growth, but also caused skin rash, diarrhea, and weight loss in mice (Extended Data Fig. 16b, c). In sum, these data suggest αNRR3 may be a safe and effective systemic therapy to treat meningiomas that are resistant to standard interventions.

## Methods

### Inclusion and ethics

This study complied with all relevant ethical regulations and was approved by the UCSF Institutional Review Board (13-12587, 17-22324, 17-23196, and 18-24633). As part of routine clinical practice at UCSF, all human patients who were included in this study signed a written waiver of informed consent to contribute deidentified data to research projects. As part of routine clinical practice at the University of California Davis, all owners of dog meningioma patients who were included in this study signed a written waiver of informed consent to contribute deidentified data to research projects. This study was approved by the UCSF Institutional Animal Care and Use Committee (AN191840), and all experiments complied with relevant ethical regulations.

### Single-cell RNA sequencing

Single cells were isolated from fresh tumor or tumor-adjacent dura samples from human or dog meningiomas, or from meningioma xenografts, as previously described^11^. Single-cell suspensions were processed for single-cell RNA sequencing using the Chromium Single Cell 3’ GEM, Library & Gel Bead Kit v3.1 (1000121, 10x Genomics) and a 10x Chromium controller, using the manufacturer recommended default protocol and settings at a target cell recovery of 5,000 cells per sample. Libraries were sequenced on an Illumina NovaSeq 6000, targeting >50,000 reads per cell, at the UCSF Center for Advanced Technology. Library demultiplexing, read alignment, identification of empty droplets, and UMI quantification were performed using CellRanger (https://github.com/10xGenomics/cellranger). Cells were filtered based on the number of unique genes and single-cell UMI count data were preprocessed in R with the Seurat^61,62^ package (v4.3.0) using the sctransform workflow^63^. Dimensionality reduction was performed using PC analysis. Uniform manifold approximation and projection (UMAP) and Louvain clustering were performed on the reduction data, followed by marker identification and differential gene expression.

Clusters were defined using a combination of automated cell type classification^26^, cell signature gene sets^27^, cell cycle analysis, and differentially expressed cluster marker genes. The ScType R package was used for automated cell type classification, with default parameters and package-provided marker genes specific to each cell type^26^. Gene set enrichment analysis was performed on clusters using cell type signature gene sets from MSigDB (https://www.gsea-msigdb.org/gsea/msigdb) with the fgsea R package (Bioconductor v3.16). Cell cycle phases of individual cells were assigned with the ‘CellCycleScoring’ function in Seurat, using single-cell cell cycle marker genes^64^.

Human meningioma single-cell samples were aligned to the GRCh38 human reference genome; filtered to cells with greater than 250 unique genes, less than 7,500 unique genes, and less than 25% of reads attributed to mitochondrial transcripts; scaled based on regression of UMI count and percentage of reads attributed to mitochondrial genes per cell; and corrected for batch effects using Harmony^25^. Parameters for downstream analysis were a minimum distance metric of 0.4 for UMAP, resolution of 0.2 for Louvain clustering, and a minimum difference in fraction of detection of 0.4 and a minimum log-fold change of 0.5 for marker identification. All human meningiomas analyzed using single-cell RNA sequencing in this study had DNA methylation profiles classifying as Immune-enriched or Hypermitotic^11^, or had biallelic inactivation of *NF2* including loss of at least 1 copy of chromosome 22q from targeted next-generation DNA sequencing^65^.

Dog meningioma single-cell samples were aligned to the ROSCfam1.0 canine reference genome; filtered to cells with greater than 1,000 unique genes and less than 6,500 unique genes; scaled based on regression of UMI count; and corrected for batch effects using Harmony^25^. Parameters for downstream analysis were a minimum distance metric of 0.2 for UMAP, resolution of 0.2 for Louvain clustering, and a minimum difference in fraction of detection of 0.25 and a minimum log-fold change of 0.8 for marker identification.

Meningioma xenograft single-cell samples were aligned to a multi-species reference genome comprised of the GRCh37 human reference genome and the GRCm38 mouse reference genome. Cells were classified as human or mouse cells based on the percentage of UMIs aligning to each genome and the distribution of those percentages. Cells with >97% of UMIs aligning to the human genome were classified as human cells, while cells with >75% of UMIs aligning to the mouse genome were classified as mouse cells. Human and mouse cells were analyzed independently after alignment.

Meningioma xenograft human tumor cells were filtered to cells with greater than 200 unique genes, less than 9,000 unique genes, and less than 20% of reads attributed to mitochondrial transcripts; and scaled based on regression of UMI count and percentage of reads attributed to mitochondrial genes per cell. Parameters for downstream analysis were a minimum distance metric of 0.1 for UMAP, resolution of 0.2 for Louvain clustering, and a minimum difference in fraction of detection of 0.3 and a minimum log-fold change of 0.25 for marker identification.

Meningioma xenograft mouse microenvironment cells were filtered to cells with greater than 250 unique genes, less than 7,500 unique genes, and less than 5% of reads attributed to mitochondrial transcripts; and scaled based on regression of UMI count and percentage of reads attributed to mitochondrial genes per cell. Parameters for downstream analysis were a minimum distance metric of 0.2 for UMAP, resolution of 0.2 for Louvain clustering, and a minimum difference in fraction of detection of 0.5 and a minimum log-fold change of 0.5 for marker identification.

### Single-cell RNA sequencing analysis

Single human meningioma cells were classified as tumor or non-tumor cells based on copy number loss of chromosome 22q. All human meningiomas analyzed using single-cell RNA sequencing in this study had copy number loss of chromosome 22q from DNA methylation profiling or targeted next-generation DNA sequencing^65^. The presence or absence of copy number variants in individual cells was assessed using the CONICSmat R package (v1.0)^28^. Briefly, a two-component Gaussian mixture model was fit to the average expression values of genes on chromosome 22q across all cells assessed. The command ‘plotAll’ from CONICSmat was run with the parameters ‘repetitions=100, postProb=0.8’. Cells with a posterior probability less than 0.2 were identified as tumor, while cells with a posterior probability greater than 0.8 were identified as normal.

The cell-cell communication network for the Notch signaling pathway was inferred and visualized using the CellChat R package (v1.5.0)^35^. Briefly, differentially expressed signaling genes were identified, noise was mitigated by calculating the ensemble average expression, intercellular communication probability was calculated by modeling ligand-receptor interactions using the law of mass action, and statistically significant communications were identified. The command ‘computeCommunProb’ from CellChat was run with the parameters ‘raw.use=FALSE, nboot=20’. All other commands were run with default parameters.

Human meningioma single-cell cluster identities were projected onto reduced dimensionality clusters of dog meningioma cells using the commands ‘FindTransferAnchors’ and ‘TransferData’ from Seurat. The parameter ‘normalization.method = ”SCT”’ was used for ‘FindTransferAnchors’ and defaults were used for all other parameters for both commands.

### Bulk RNA-sequencing analysis

Human meningioma bulk RNA sequencing data were generated and analyzed as previously described^11,13^. In brief, RNA was extracted from frozen meningiomas, and library preparation was performed using the TruSeq Standard mRNA Kit (20020595, Illumina) 50 bp single-end or 150 bp paired-end reads that were sequenced on an Illumina NovaSeq 6000 to a mean of 20 million reads per sample. Analysis was performed using a pipeline comprised of FastQC for quality control, and Kallisto for reading pseudoalignment and transcript abundance quantification using the default settings (v0.46.2).

Gene set enrichment analysis (GSEA v4.3.2) was performed to identify differentially expressed pathways distinguishing recurrent from primary meningiomas. Gene rank scores were calculated using the formula sign(log_2_ fold-change) × −log10(p-value). Pathways were defined using the gene set file Human_GOBP_AllPathways_no_GO_iea_December_01_2022_symbol.gmt, which is maintained by the Bader laboratory. Gene set size was limited to range between 15 and 500, and positive and negative enrichment files were generated using 2000 permutations. The EnrichmentMap App (v3.3.4) in Cytoscape (v3.7.2) was used to visualize the results of pathway analysis. Nodes with p≤0.01 and FDR≤0.01, and nodes sharing gene overlaps with Jaccard + Overlap Combined (JOC) threshold of 0.375 were connected by blue lines (edges) to generate network maps. Clusters of related pathways were identified and annotated using the AutoAnnotate app (v1.3.5) in Cytoscape that uses a Markov Cluster algorithm to connect pathways by shared keywords in the description of each pathway. The resulting groups of pathways were designated as the consensus pathways in each circle.

### Histology, immunohistochemistry, immunofluorescence, and microscopy

For adult human tissue samples, deparaffinization and rehydration of 5µm formalin-fixed, paraffin-embedded (FFPE) tissue sections and H&E staining were performed using standard procedures. Immunostaining was performed on an automated Ventana Discovery Ultra staining system and detection was performed with Multimer HRP (Ventana Medical Systems) followed by fluorescent detection (DISCOVERY Rhodamine and CY5) or DAB. Immunostaining for NOTCH3 was performed using mouse monoclonal NOTCH3/N3ECD primary antibody (MABC594, Millipore Sigma, 1:25-1:100) with incubation for 32min following CC1 antigen retrieval for 32min. For dual staining, primary antibody incubations were carried out serially with inclusion of positive, negative, and single antibody controls. Following staining for NOTCH3/N3ECD, tissue sections were stained with primary antibodies recognizing CD34 (CBL496, Millipore Sigma, mouse monoclonal, 1:300) for 2h, SMA (ab7817, Abcam, mouse polyclonal, 1:30,000) for 32min, or VWF (A0082, Dako, rabbit polyclonal, 1:1,000) for 20min. All IF experiments were imaged on a LSM 800 confocal laser scanning microscope with Airyscan (Zeiss) and analyzed using ImageJ.

For meningioma xenograft samples, deparaffinization and rehydration of 5µm FFPE tissue sections and H&E staining were performed using standard procedures. IHC was performed on an automated Ventana Discovery Ultra staining system using primary antibodies recognizing Ki-67 (M7240, DAKO, mouse monoclonal, clone MIB1, 1:50) for 30min, CD34 (NCL-L-END, LEICA, mouse monoclonal, clone QBEnd/10, undiluted) for 15min, or ERG (790-4576, Ventana, rabbit monoclonal, clone EPR3864, undiluted) for 32min. All histological and IHC experiments were imaged on a BX43 light microscope (Olympus) and analyzed using the Olympus cellSens Standard Imaging Software package.

For IF of the developing human brain, de-identified tissue samples were collected with previous patient consent in strict observance of legal and institutional ethical regulations. Autopsy consent and all protocols were approved by the UCSF Human Gamete, Embryo, and Stem Cell Research Committee. All cases were determined by chromosomal analysis, physical examination, and/or pathological analysis to be control tissues, which indicates that they were absent of neurological disease. Brains were cut into ∼1.5cm coronal or sagittal blocks, fixed in 4% paraformaldehyde for 2 days, and then cryoprotected in a 30% sucrose solution. Blocks were cut into 30µm sections on a cryostat, mounted on glass slides for IF, and stored at -80°C. Frozen slides were moved from -80°C to 4°C the night prior to staining and then to the lab bench for 2h before beginning the immunostaining protocol. Slides were washed once with 1X PBS for 5 minutes, then once with 1X TBS for 5min before blocking with TBS++++ (goat serum, BSA, albumin, glycine, and triton X in TBS) for 1h. Primary antibodies recognizing PDGFRB (AF385, R&D Systems, 1:200), CD34 (AF7227, R&D Systems, 1:200), or NOTCH3/N3ECD as described above were used with overnight incubation at room temperature at 1:200 dilutions in TBS++++. The following day, three 1x TBS washes were performed before incubating with secondary antibodies in TBS++++ for 2h. After three additional TBS washes, DAPI (62248, ThermoFisher Scientific) was added, and the slides were mounted.

Immunofluorescence of meningioma cell lines were performed on cover glass slips in culture. Cells were fixed in 4% PFA in PBS for 8min, washed in PBS, blocked for 30min in 5% donkey serum and 0.1% Triton-X100 in PBS. Cells were staining with PTPRZ1 (sc-33664, Santa Cruz Biotechnology, 1:1000) overnight at 4°C and subsequently labeled with rabbit Alexa Fluor secondary antibody (A21206, ThermoFischer Scientific, 1:1000) and Hoechst 33342 to mark DNA for 1h at room temperature prior to mounting and imaging. Meningioma cells were imaged on a LSM 800 confocal laser scanning microscope with Airyscan (Zeiss) and analyzed using ImageJ.

### RNAScope and microscopy

The RNAScope Multiplex Fluorescent V2 assay (32310, ACDBio) was performed according to the manufacturer’s protocol. Briefly, 5µm FFPE meningioma sections were incubated with hydrogen peroxide to inhibit endogenous peroxidase, followed by processing for target retrieval and treatment with ProteasePlus. Meningioma sections were subsequently incubated with RNA probes for *NOTCH3* (558991-C2, Hs-NOTCH3-C2) and *NF2* (1037481-C, Hs-NF2-C1), followed by revelation and amplification steps. Meningioma sections were blocked (5% normal donkey serum, 1X Animal Free blocking, 0.3% Triton X-100) for 1 hour at room temperature and incubated with primary antibody against VE-cadherin (AF938, R&D Systems, 1:100) overnight at 4°C. The next day, meningioma sections were incubated with secondary antibodies for 1 hour at room temperature and counterstained with DAPI (62248, ThermoFisher Scientific). Meningioma sections were imaged on a LSM 800 confocal laser scanning microscope with Airyscan (Zeiss) and analyzed using ImageJ.

### Multiplexed sequential immunofluorescence (seqIF) and microscopy

Automated multiplexed seqIF staining and imaging was performed on FFPE sections at Northwestern University using the COMET platform (Lunaphore Technologies). The multiplexed panel was comprised of 29 antibodies (Supplementary Table 4). The 29-plex protocol was generated using the COMET Control Software, and reagents were loaded onto the COMET device to perform seqIF. All antibodies were validated using conventional IHC and/or IF staining in conjunction with corresponding fluorophores and DAPI (62248, ThermoFisher Scientific). For optimal concentration and best signal-to-noise ratio, all antibodies were tested at 3 different dilutions: starting with the manufacturer-recommended dilution (MRD), MRD/2, and MRD/4. Secondary Alexa fluorophore 555 (A32727, ThermoFisher Scientific) and Alexa fluorophore 647 (A32733, ThermoFisher Scientific) were used at 1:200 or 1:400 dilutions, respectively. The optimizations and full runs of the multiplexed panel were executed using the seqIF technology integrated in the Lunaphore COMET platform (characterization 2 and 3 protocols, and seqIF protocols, respectively). The seqIF workflow was parallelized on a maximum of 4 slides, with automated cycles of iterative staining of 2 antibodies at a time, followed by imaging, and elution of the primary and secondary antibodies, with no sample manipulation during the entire workflow. All reagents were diluted in Multistaining Buffer (BU06, Lunaphore Technologies). The elution step lasted *2*min for each cycle and was performed with Elution Buffer (BU07-L, Lunaphore Technologies) at 37°C. Quenching lasted for 30sec and was performed with Quenching Buffer (BU08-L, Lunaphore Technologies). Incubation time was set at 4min for all primary antibodies, except for the p16 antibody at 8min, and secondary antibodies at 2min. Imaging was performed with Imaging Buffer (BU09, Lunaphore Technologies) with exposure times set for 400 ms for the TRITC channel, 200ms for the Cy5 channel, and 80ms for the DAPI channel. Imaging was performed with Imaging Buffer (BU09, Lunaphore Technologies) with exposure times set at 4min for all primary antibodies, except P16 antibody at 8min, and secondary antibodies at 2min. Imaging was performed with an integrated epifluorescent microscope at 20x magnification. Image registration was performed immediately after concluding the staining and imaging procedures by COMET Control Software. Each seqIF protocol resulted in a multi-stack OME-TIFF file where the imaging outputs from each cycle were stitched and aligned. COMET OME-TIFF files contain a DAPI image, intrinsic tissue autofluorescence in TRITC and Cy5 channels, and a single fluorescent layer per marker. Markers were subsequently pseudocolored for visualization of multiplexed antibodies.

### Mouse genetic models

Notch3^tm1.1(cre/ERT2)Sat^ (*Notch3-Cre*^ERT2^) mice were obtained from the Sweet-Cordero Lab at UCSF. Ptgds^tm1.1(cre)Gvn^ (*Ptgds-Cre*) mice were obtained from Riken. Gt(ROSA)26Sor^tm4(ACTB-tdTomato,-EGFP)Luo^ (*ROSA*^mT/mG^) mice were obtained from The Jackson Laboratory. Gt(ROSA)26Sor^tm1(HBEGF)Awai/J^ (*ROSA*^iDTR^) mice were obtained from The Jackson Laboratory. FVB.129P2-Nf2^tm2Gth^ (*Nf2*^fl/fl^) mice were obtained from Riken. All mouse genetic experiments were performed on the C57BL/6J background. Mice were intercrossed to generate *Notch3-Cre*^ERT2(+/WT)^ *ROSA*^mT/mG^ mice, *Ptgds-Cre*^+/WT^ *ROSA*^mT/mG^ mice, *Notch3-Cre*^ERT2(+/WT)^ *Nf2*^fl/fl^ mice, and *Notch3-Cre*^ERT2(+/WT)^ *ROSA*^iDTR^ mice. Recombination of either the *Nf2* or *ROSA* locus was induced using intraperitoneal injection of 75mg/kg of tamoxifen (T5648, Sigma-Aldrich) dissolved in corn oil. For *in utero* recombination, pregnant dams were injected with tamoxifen once at E16.5 and their subsequent litters were genotyped and euthanized at the indicated timepoints. For postnatal recombination, mice were injected daily 5 times with tamoxifen and euthanized at the indicated timepoints.

Mouse skullcaps for meningeal lineage tracing experiments and intact skulls for tumorigenesis experiments were processed as previously described^66^. In brief, mice were perfused with 4°C 4% paraformaldehyde in 80nM Pipes, pH 6.8, 5mM EGTA, and 2mM MgCl_2_. Muscle was dissected and removed, and samples were rotated overnight at 4°C in perfusion solution. Samples were washed three times for 5min in PBS and decalcified for 3 days rotating at 4°C in 20% EDTA, pH 7.4, in PBS. For meningioma tumorigenesis experiments, whole skulls were subsequently embedded in 5% low-melt agarose (Precisionary) and cut into 300µm sections on a Vibratome (VT1000S, Leica). Sections were stained for 15s in Mayer’s Hematoxylin solution (MHS16, Sigma-Aldrich), washed three times for 5min in distilled water, incubated for 1min in PBS to blue nuclei, washed for 5min in distilled water, stained for 30s in 0.5% Eosin Y (1099884, Sigma-Aldrich), washed two times for 5min in 70% ethanol, and mounted on slides using Vectashield Antifade Mounting Media (H-1000-10, Vector Laboratories).

### Mouse xenograft models and treatments

Xenograft experiments were performed by implanting CH-157MN or IOMM-Lee cells into the flank of 5-to 6-week-old female NU/NU mice (Harlan Sprague Dawley) as previously described^11^. Tumor initiating capacity was defined by the development of sustained, iteratively increasing subcutaneous growth at injection sites. Radiotherapy treatments were performed using a Precision X-RAD 320 Cabinet Irradiator with normal operating settings to deliver 2Gy of ionizing radiation on each of 5 consecutive days. NOTCH3 negative regulatory region neutralizing antibody treatments (αNRR3) and NOTCH1 negative regulatory region neutralizing antibody (αNRR1) treatments using murine antibodies from Genentech were performed as previously described^7,8^, with bi-weekly IP injection of 20mg/kg αNRR3, 10mg/kg αNRR1 (dose-reduced due to gastrointestinal and cutaneous toxicity leading to weight loss), or 20mg/kg IgG2a isotype control (BE0085, Bio X-cell). γ-secretase inhibitor treatments were performed using LY-411575 (SML0649, Millipore Sigma) as previously described^47^, with daily 20µM/kg IP injections in 0.5% methylcellulose and 0.1% Tween 80 in 1xPBS. For Kaplan-Meier survival analyses, events were recorded when tumors reached the protocol-defined size of 2000 mm^3^, mice developed mobility or physiological impairment from tumor burden, or mice lost >15% of body weight due to treatment-associated toxicity.

### Cell culture

HEK293T (CRL-3216, ATCC), IOMM-Lee^51^, or CH157-MN^50^ cells were cultured in Dulbecco’s modified Eagle medium (DMEM) (11960069, Life Technologies) supplemented with 10% fetal bovine serum (FBS), 1x GlutaMAX (35050-061, ThermoFischer Scientific) and 1x Penicillin/Streptomycin (15140122, Life Technologies).

To generate cell lines overexpressing NOTCH3 ICD, pLVX-Puro plasmid containing pCMV6-NOTCH3ICD was generated. Lentiviral particles were produced by transfecting HEK293T cells with standard packaging vectors using the *Trans*IT-Lenti Trasfection Reagent (6605, Mirus). CH157-MN cells were stably transduced with lentiviral particles to generate CH-157MN NOTCH3^ICD^ or empty pLVX vector (EV) cells. Successfully transduced cells were isolated using Puromycin selection, and NOTCH3 overexpression was confirmed using QPCR.

For clonogenic assays, 250 cells were seeded in triplicate for each condition in a 6-well plate and grown for 10 days in standard culture media. Cells were fixed in methanol for 30min, washed and stained in 0.01% crystal violet (C6158, Sigma-Aldrich) for 3hours, washed 3 times in distilled water, and dried overnight. Cells were imaged on a Stemi 508 stereo microscope (Zeiss) and clonogenic area was quantified in ImageJ.

### Immunoblotting

Meningioma cells for immunoblotting were lysed in 1% SDS in 100mM pH 6.8 Tris-HCL containing protease and phosphatase inhibitor (A32961, Thermo Scientific), vortexed at maximum speed for 1min, rocked at 4°C for 5min, and centrifuged at 4°C for 15min at 15000xg. Supernatant protein quantification was performed using BCA assays (23225, Pierce). 20μg of lysate from each cell line was boiled for 15 min in Laemmli reducing buffer. Proteins were separated on 4-15% TGX precast gels (5671084, Bio-Rad), and transferred onto ImmunBlot PVDF membrane (1620177, Bio-Rad). Membranes were blocked in 5% TBST-milk, incubated in primary antibodies, washed, and incubated in secondary antibodies. Membranes were subjected to immunoblot analysis using Pierce ECL substrate (32209, Thermo Fischer Scientific). Primary antibodies recognizing NOTCH3 (5276, Cell Signaling, 1:1000) or GAPDH (8245, Abcam, 1:5000) and secondary antibodies recognizing mouse (7076, Cell Signaling, 1:2000) or rabbit (7074, Cell Signaling, 1:2000) epitope were used.

### Quantitative reverse-transcriptase polymerase chain reaction

RNA was extracted from cultured meningioma cells using the RNeasy Mini Kit (74106, QIAGEN) according to manufacturer’s instructions. cDNA was synthesized from extracted RNA using the iScript cDNA Synthesis Kit (1708891, Bio-Rad). Real-time QPCR was performed using PowerUp SYBR Green Master Mix (A25918, Thermo Fisher Scientific) on a QuantStudio 6 Flex Real Time PCR system (Life Technologies) using forward and reverse primers (*NOTCH3-F* CGTGGCTTCTTTCTACTGTGC, *NOTCH3-R* CGTTCACCGGATTTGTGTCAC, *HEY1-F* GTTCGGCTCTAGGTTCCATGT, *HEY1-R* CGTCGGCGCTTCTCAATTATT, *GAPDH-F* GTCTCCTCTGACTTCAACAGCG, *GAPDH-R* ACCACCCTGTTGCTGTAGCCAA). QPCR data were analyzed using the ΔΔCt method relative to *GAPDH* expression.

### Statistics

All experiments were performed with independent biological replicates and repeated, and statistics were derived from biological replicates. Biological replicates are indicated in each figure panel or figure legend. No statistical methods were used to predetermine sample sizes, but sample sizes in this study are similar or larger to those reported in previous publications. Data distribution was assumed to be normal, but this was not formally tested. Investigators were blinded to conditions during clinical data collection and analysis of mechanistic or functional studies. Bioinformatic analyses were performed blind to clinical features, outcomes, or molecular characteristics. The clinical samples used in this study were retrospective and nonrandomized with no intervention, and all samples were interrogated equally. Thus, controlling for covariates among clinical samples was not relevant. Cells and animals were randomized to experimental conditions. No clinical, molecular, or cellular data points were excluded from the analyses. Lines represent means, and error bars represent standard error of the means. Results were compared using Student’s t-tests and ANOVA, which are indicated in figure panels or figure legends alongside approaches used to adjust for multiple comparisons. In general, statistical significance is shown using asterisks (*p≤0.05, **p≤0.01, ***p≤0.0001), but exact p*-*values are provided in figure panels or figure legends when possible.

### Reporting summary

Further information on research design is available in the Nature Research Reporting Summary linked to this article.

## Data availability

Single-cell RNA sequencing data of new samples (n=1 human meningioma sample, n=3 canine meningioma samples, n=2 canine dura samples, n=23 meningioma xenograft samples) that are reported in this manuscript will be deposited in the NCBI Gene Expression Omnibus upon provisional acceptance and are available during the review process upon request. Additional single-cell RNA sequencing data from previously reported human meningiomas (n=4 meningioma samples, n=1 dura sample) are available under the accession GSE183655 (https://www.ncbi.nlm.nih.gov/geo/query/acc.cgi?acc=GSE183655). RNA sequencing and DNA methylation profiling data of human meningioma samples (n=502) are available under the accessions GSE183656 (https://www.ncbi.nlm.nih.gov/geo/query/acc.cgi?acc=GSE183656), GSE212666 (https://www.ncbi.nlm.nih.gov/geo/query/acc.cgi?acc=GSE212666), GSE183653 (https://www.ncbi.nlm.nih.gov/geo/query/acc.cgi?acc=GSE183653), and GSE101638 (https://www.ncbi.nlm.nih.gov/geo/query/acc.cgi). The publicly available GRCh38 (hg38, https://www.ncbi.nlm.nih.gov/assembly/GCF_000001405.39/), GRCh37 (hg19, https://www.ncbi.nlm.nih.gov/assembly/GCF_000001405.25/), GRCm38 (mm10, https://www.ncbi.nlm.nih.gov/assembly/GCF_000001635.20/), and ROS_Cfam_1.0 datasets (https://www.ncbi.nlm.nih.gov/assembly/GCF_014441545.1/) were used in this study. Source data are provided with this paper.

## Code availability

The open-source software, tools, and packages used for data analysis in this study, as well as the version of each program, were ImageJ (v2.1.0), CellRanger (v6.1.2 and v7.1.0), kallisto (v0.46.2), GSEA (v4.3.2), EnrichmentMap (v3.3.4), Cytoscape (v3.7.2), AutoAnnotate (v1.3.5), R (v4.2.3), Seurat R package (v4.3.0), Harmony R package (v0.1.1), ScType R package (v1), CONICSmat R package (v1.0), CellChat R package (v1.5.0), fgsea (Bioconductor v3.16), and DESeq2 (Bioconductor v3.16). No software was used for data collection. No custom algorithms or code were used.

## Acknowledgements

The authors thank Mark Youngblood, Julie Siegenthaler, Sten Linnarsson, Elin Vinsland, Chris Siebel, Fred De Sauvage, Alejandro Sweet-Cordero, Kieren Marini, Kyla Foster, and members of the Raleigh lab for reagents, guidance, and edits, Beverly Sturges and Chai Fei Li for providing dog meningioma samples, Shai and the staff of the UCSF Brain Tumor Center Biorepository and Pathology Core, Eric Chow and the staff of the UCSF Center for Advanced Technology, and Ken Probst and Noel Sirivansanti for Illustrations. Lunaphore COMET multiplexed seqIF was enabled by a gift from the Stephen M. Coffman trust to the Northwestern University Malnati Brain Tumor Institute of the Lurie Cancer Center. This study was supported by the UCSF Wolfe Meningioma Program Project and the National Institutes of Health (NIH) grants F30 CA246808 and T32 GM007618 to AC; NIH grants T32 CA15102 and P50 CA097257 to C-HGL; NIH grant P50 CA097257 to JJP; NIH grant P50 CA097257 to WCC; the UCSF Wolfe Meningioma Program Project and P50 CA097257 to NAOB; NIH grants R01 NS117104, R01 NS118039, and P50 CA221747 to CMH; the UCSF Wolfe Meningioma Program Project, NIH grant F32 CA213944, and the Northwestern Medicine Malnati Brain Institute of the Lurie Cancer Center to STM; NIH grants R01 CA120813, R01 NS120547, and P50 CA221747 to ABH; and NIH grants R01 CA262311 and P50 CA097257, and the UCSF Wolfe Meningioma Program Project to DRR.

## Author contributions statement

All authors made substantial contributions to the conception or design of the study; the acquisition, analysis, or interpretation of data; or drafting or revising the manuscript. All authors approved the manuscript. All authors agree to be personally accountable for individual contributions and to ensure that questions related to the accuracy or integrity of any part of the work are appropriately investigated and resolved and the resolution documented in the literature. AC and MAC designed, performed, and analyzed the majority of the experiments and bioinformatic analyses. C-HGL performed light microscopy and analyses of human and dog meningioma histology and IHC. HN performed and analyzed multiplexed seqIF with supervision from CMH and ABH. JJP supervised meningioma histology, IHC, and IF. BP performed meningioma *in situ* hybridization and IF under supervision by OK. NZ and WCC performed bioinformatic analyses of meningiomas under supervision by AC and DRR. TJ and JOB performed bioinformatic analyses of developing and developed human cerebral vasculature under supervision by AB and EEC. NAOB, SLH-J, and STM provided human clinical data and samples. CMT and PJD provided dog clinical data and samples. DRR performed IF and fluorescence confocal microscopy, light microscopy of mouse genetic models, mouse treatments, xenograft measurements, analyses of *in vivo* data, and conceived, designed, and supervised the study.

## Tables

Not applicable

