## Extended Data Figures for "NOTCH3 drives meningioma tumorigenesis and resistance to radiotherapy"

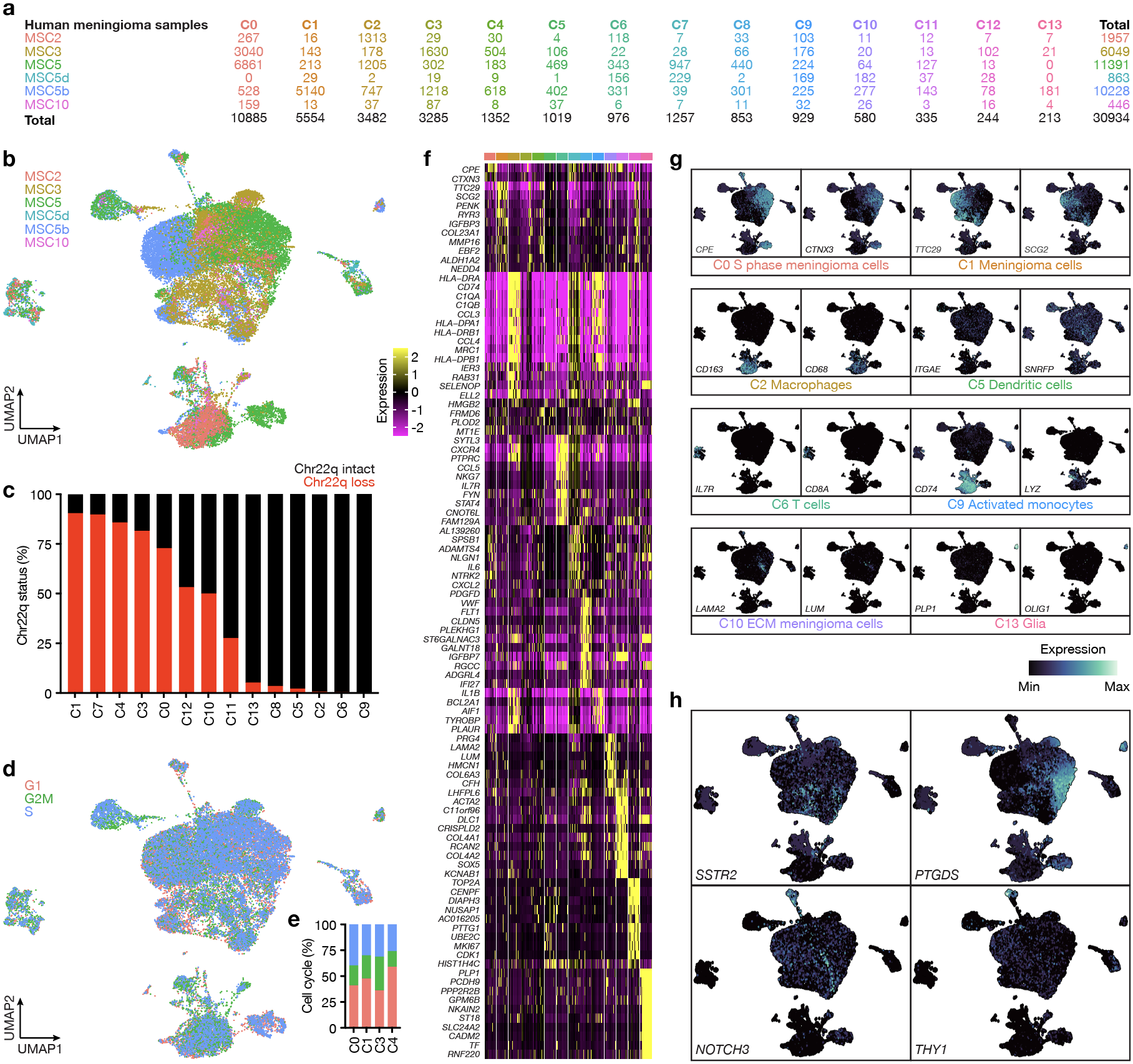
**Extended Data Fig. 1. Single-cell RNA sequencing of human meningiomas with loss of chromosome 22q. a**, Single-cell counts across UMAP clusters from human meningioma samples with loss of chromosome 22q analyzed using single-cell RNA sequencing. **b**, UMAP showing single-cell RNA sequencing of human meningioma samples shaded by sample of origin. **c**, Stacked bar plots showing the distribution of chromosome 22q loss (red) across single-cell RNA sequencing clusters from human meningioma samples. **d**, **e**, UMAP and stacked bar plot showing single-cell RNA sequencing cell cycle analysis across human meningioma samples. **f**, **g**, Heatmap and feature plots showing differentially expressed genes across UMAP clusters from human meningioma samples analyzed using single-cell RNA sequencing. Colors as in **a**. See also Supplementary Table 1. **h**, Feature plots showing expression of meningioma (*SSTR2*, *PTGDS*) or cancer stem-cell markers (*NOTCH3*, *THY1*) across UMAP clusters from human meningioma samples analyzed using single-cell RNA sequencing.

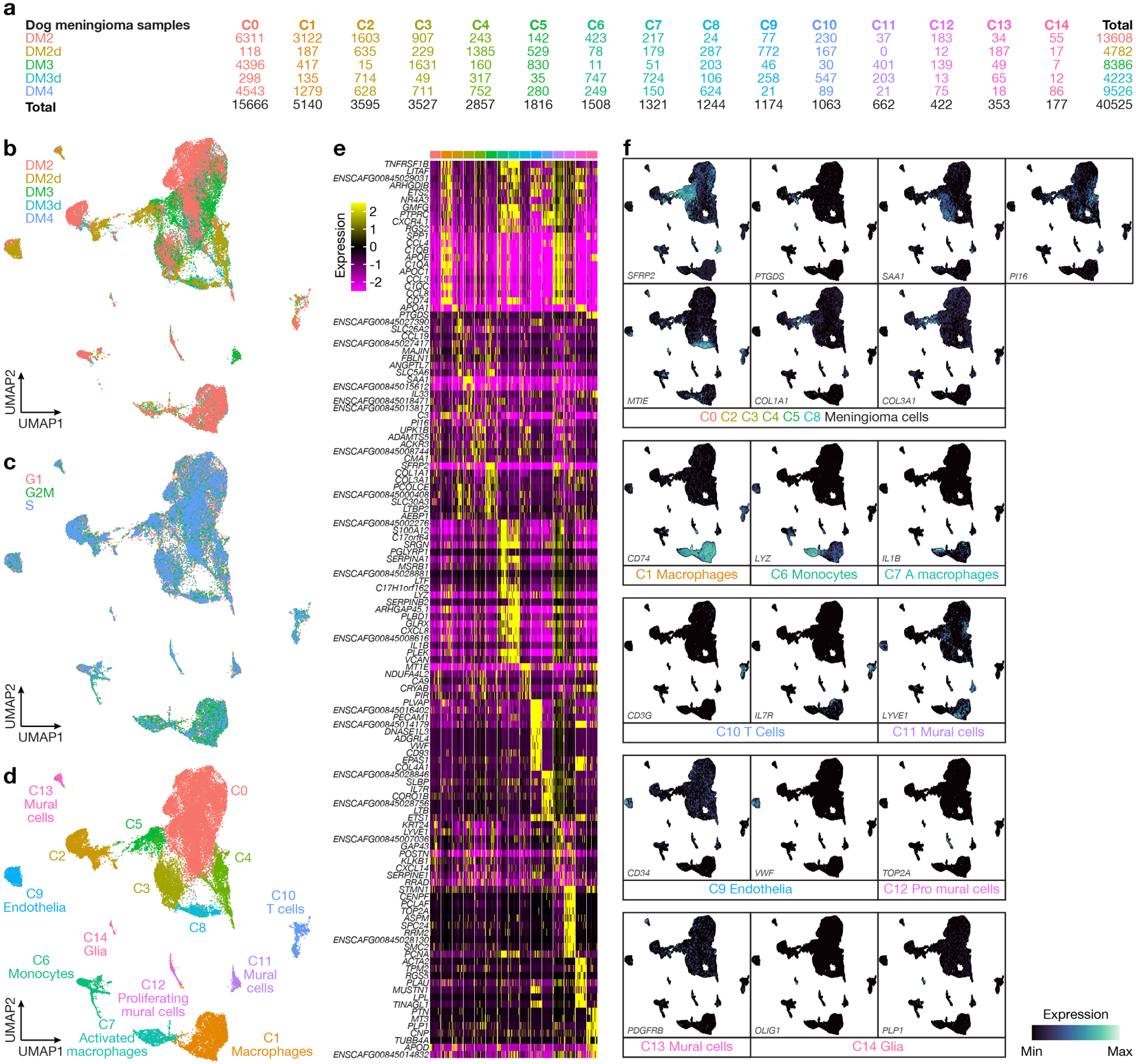
**Extended Data Fig. 2. Single-cell RNA sequencing of dog meningiomas.** **a**, Single-cell counts across UMAP clusters from dog meningioma samples analyzed using single-cell RNA sequencing. **b**, UMAP showing single-cell RNA sequencing of dog meningioma samples shaded by sample of origin. **c**, UMAP showing single-cell RNA sequencing cell cycle analysis across dog meningioma samples. **d**, Single-cell RNA sequencing UMAP of 40,525 dog meningioma transcriptomes showing tumor cell states and microenvironment cell types. **e**, **f**, Feature plots showing differentially expressed genes across UMAP clusters from dog meningioma samples analyzed using single-cell RNA sequencing. Colors as in **a**. See also Supplementary Table 2.

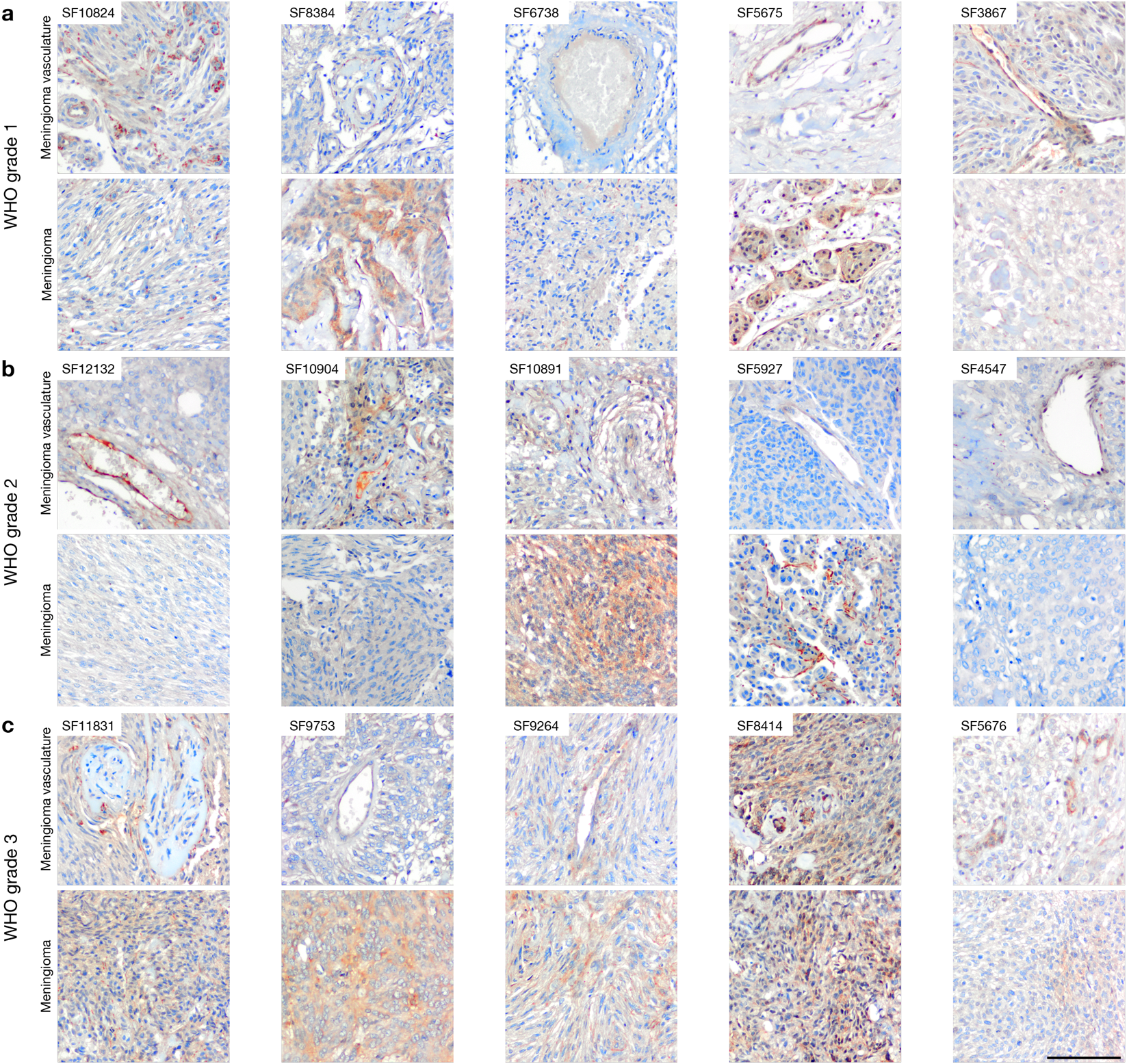
**Extended Data Fig. 3. NOTCH3 immunohistochemistry across meningioma WHO grades.** NOTCH3 expression is predominantly restricted to the perivascular niche in WHO grade 1 meningiomas in **a**, but extends beyond the perivascular niche in WHO grade 2 meningiomas in **b** and in WHO grade 3 meningiomas in **c**. SF numbers indicate individual meningiomas. WHO grade defined using histological criteria. Scale bar, 100µm.

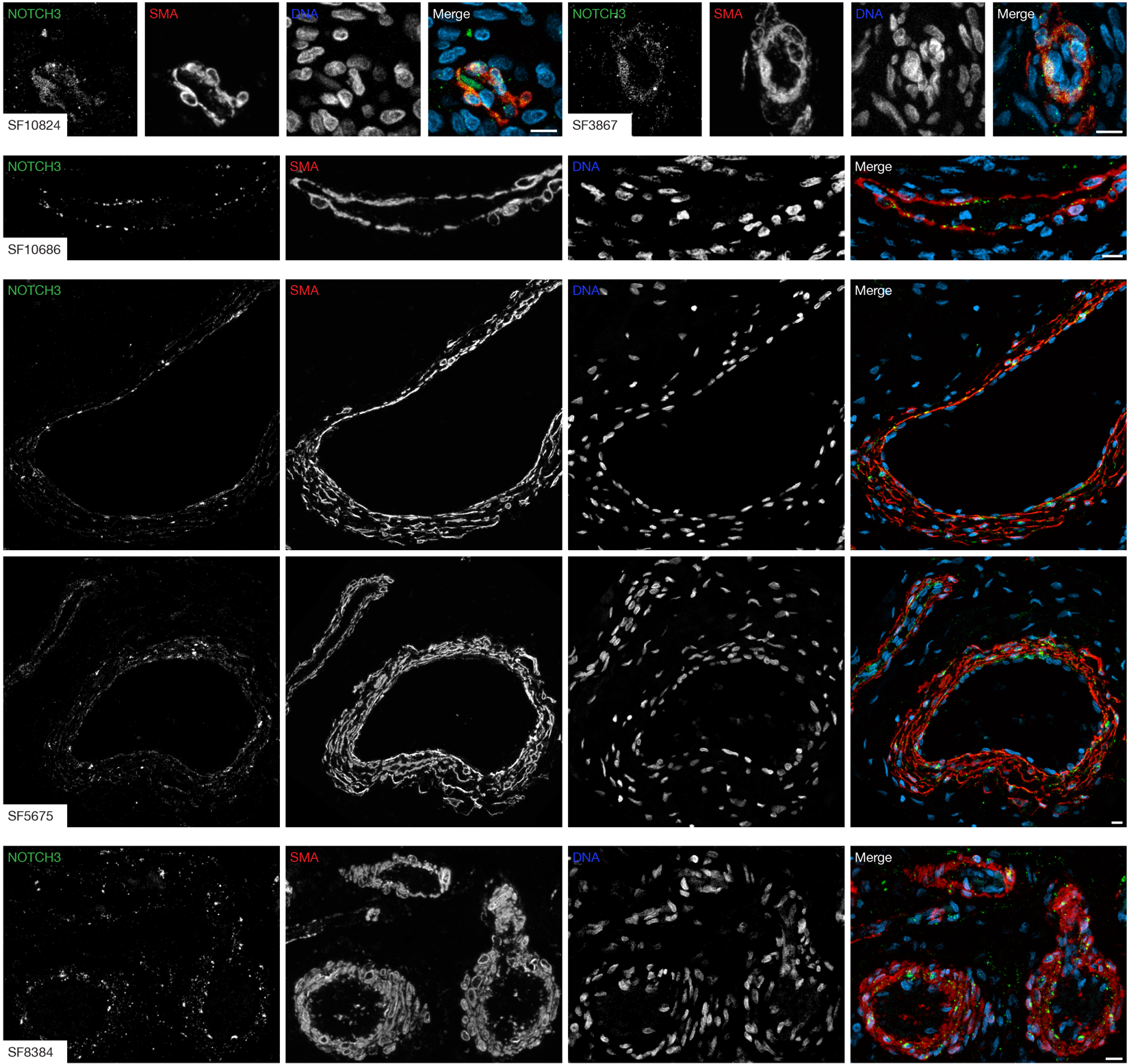

**Extended Data Fig. 4. WHO grade 1 meningioma immunofluorescence microscopy for NOTCH3 and the mural cell marker SMA.** NOTCH3 expression is restricted to the perivascular niche and colocalizes with mural cells in meningiomas with WHO grade 1 histology. SF numbers indicate individual meningiomas. WHO grade defined using histological criteria. Scale bars, 10µm.

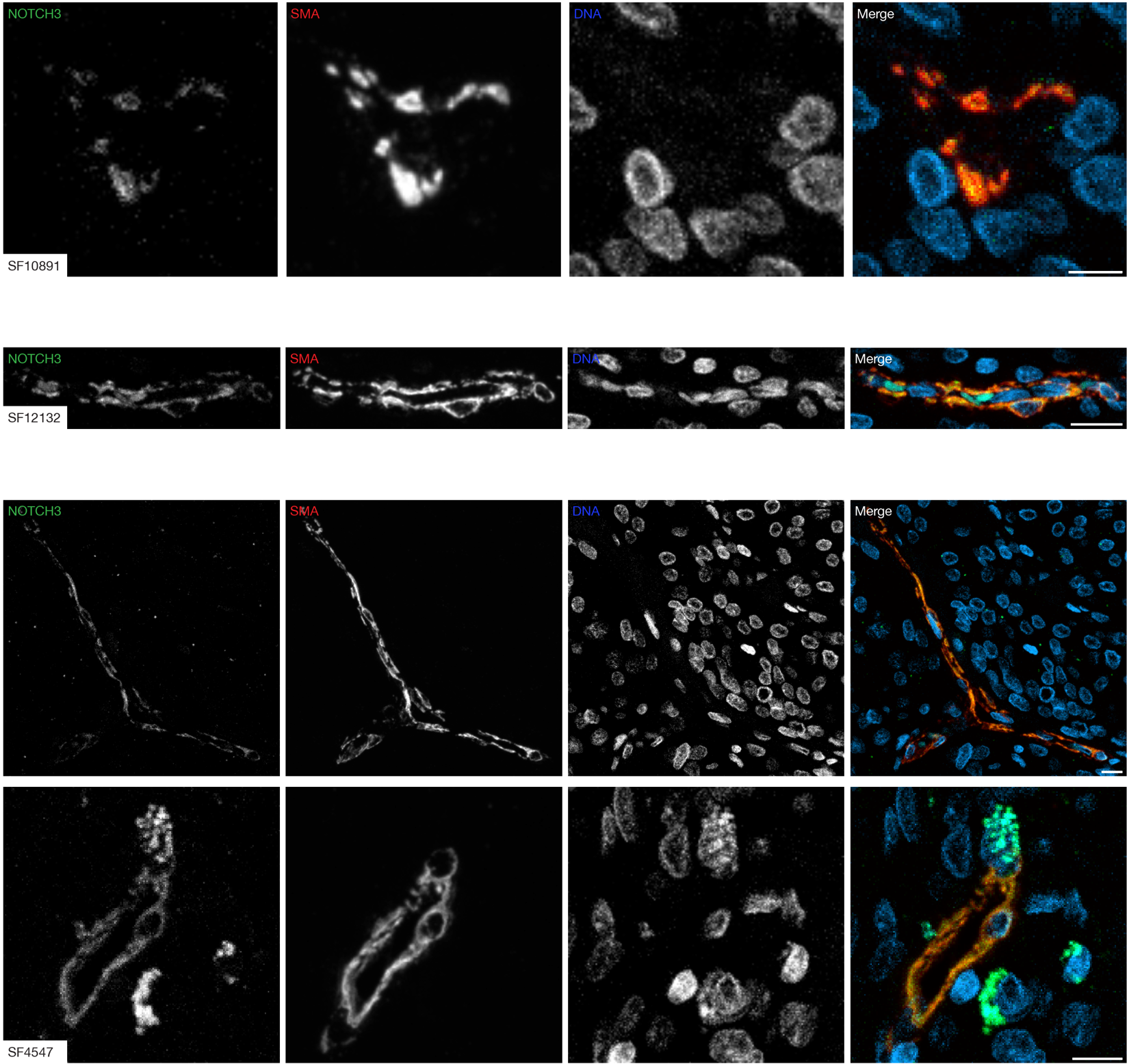
**Extended Data Fig. 5. WHO grade 2 meningioma immunofluorescence microscopy for NOTCH3 and the mural cell marker SMA.** NOTCH3 expression is primarily restricted to the perivascular niche and colocalizes with mural cells in meningiomas with WHO grade 2 histology. SF numbers indicate individual meningiomas. WHO grade defined using histological criteria. Scale bars, 10µm.

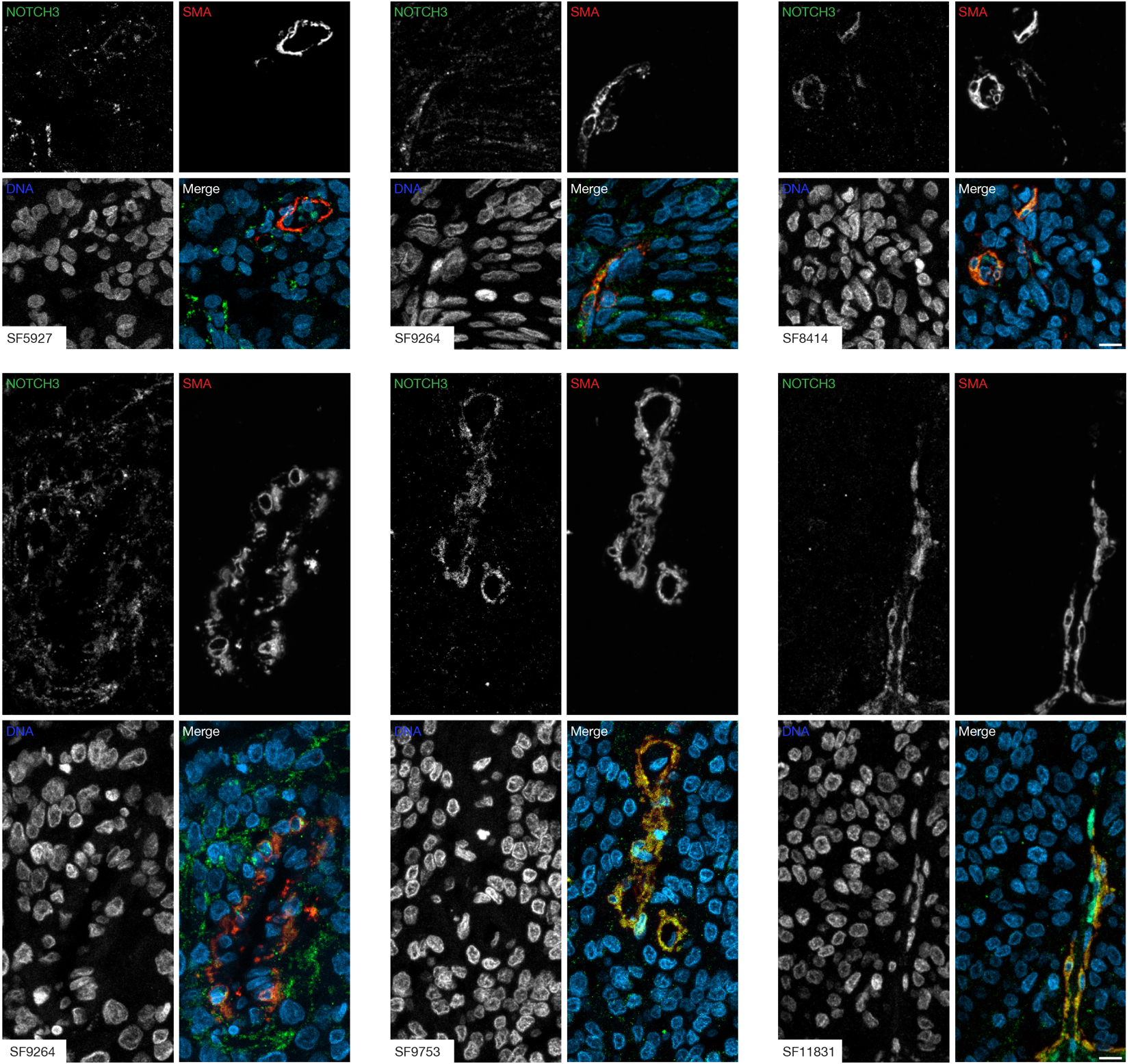
**Extended Data Fig. 6. WHO grade 3 meningioma immunofluorescence microscopy for NOTCH3 and the mural cell marker SMA.** NOTCH3 is expressed in and out of the perivascular niche, and colocalizes with mural cells in the perivascular niche, in meningiomas with WHO grade 3 histology. SF numbers indicate individual meningiomas. WHO grade defined using histological criteria. Scale bars, 10µm.

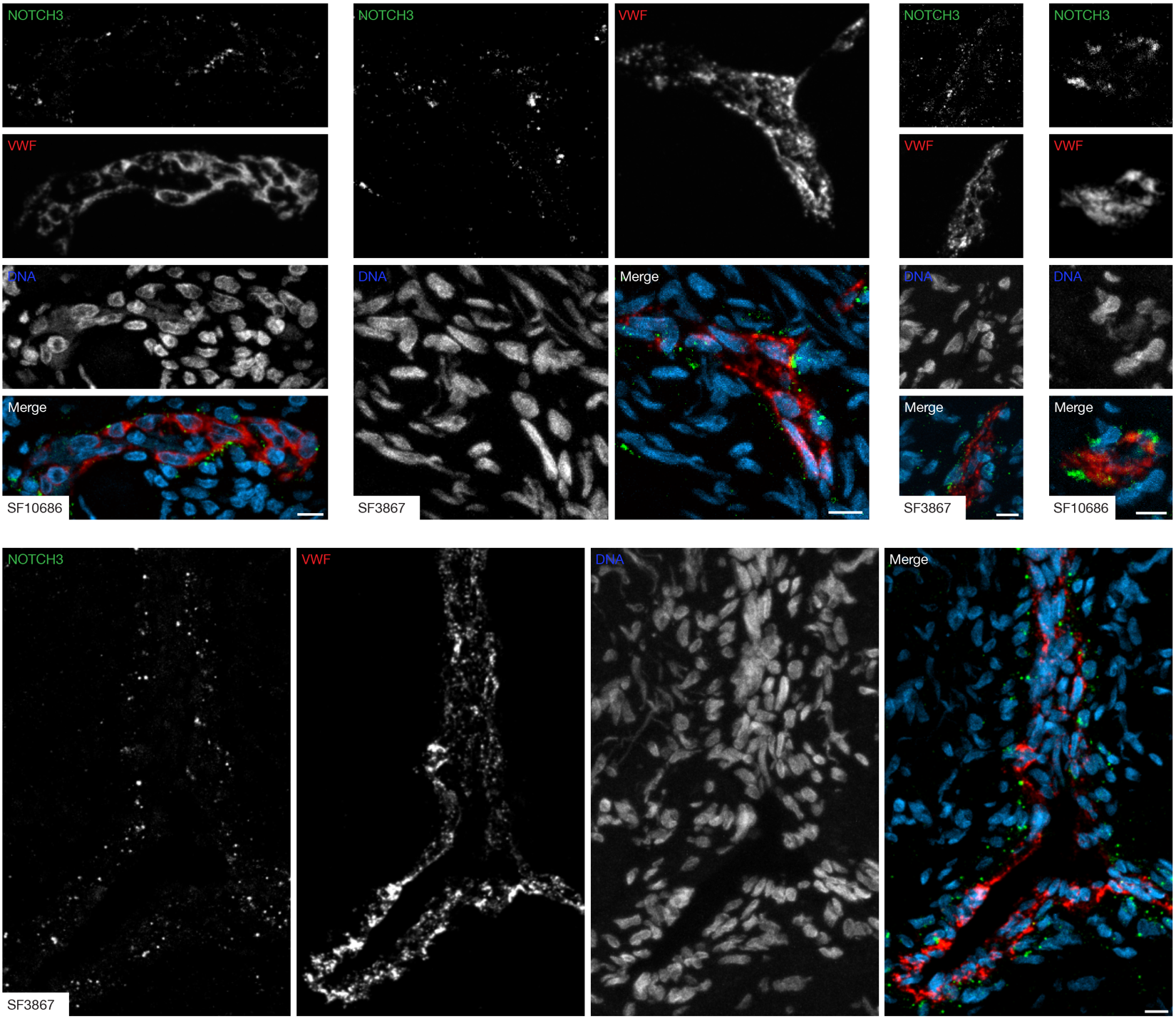
**Extended Data Fig. 7. WHO grade 1 meningioma immunofluorescence microscopy for NOTCH3 and the endothelial cell marker VWF.** NOTCH3 expression is restricted to the perivascular niche adjacent to endothelial cells in meningiomas with WHO grade 1 histology. SF numbers indicate individual meningiomas. WHO grade defined using histological criteria. Scale bars, 10µm.

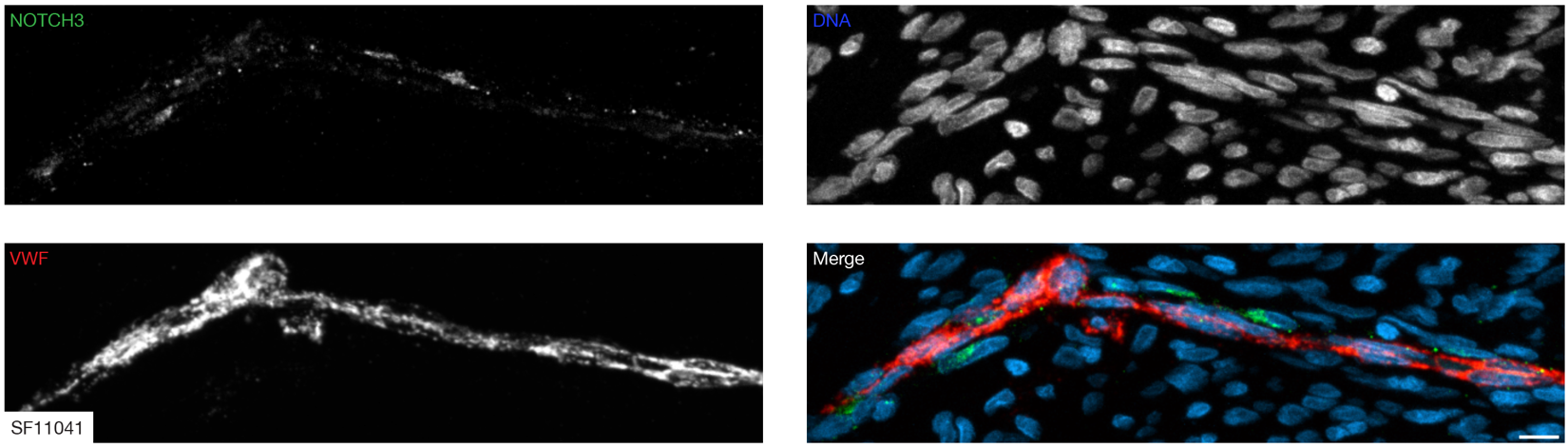
**Extended Data Fig. 8. WHO grade 2 meningioma immunofluorescence microscopy for NOTCH3 and the endothelial cell marker VWF.** NOTCH3 expression is primarily restricted to the perivascular niche adjacent to endothelial cells in meningiomas with WHO grade 2 histology. SF number indicates meningioma identifier. WHO grade defined using histological criteria. Scale bar, 10µm.

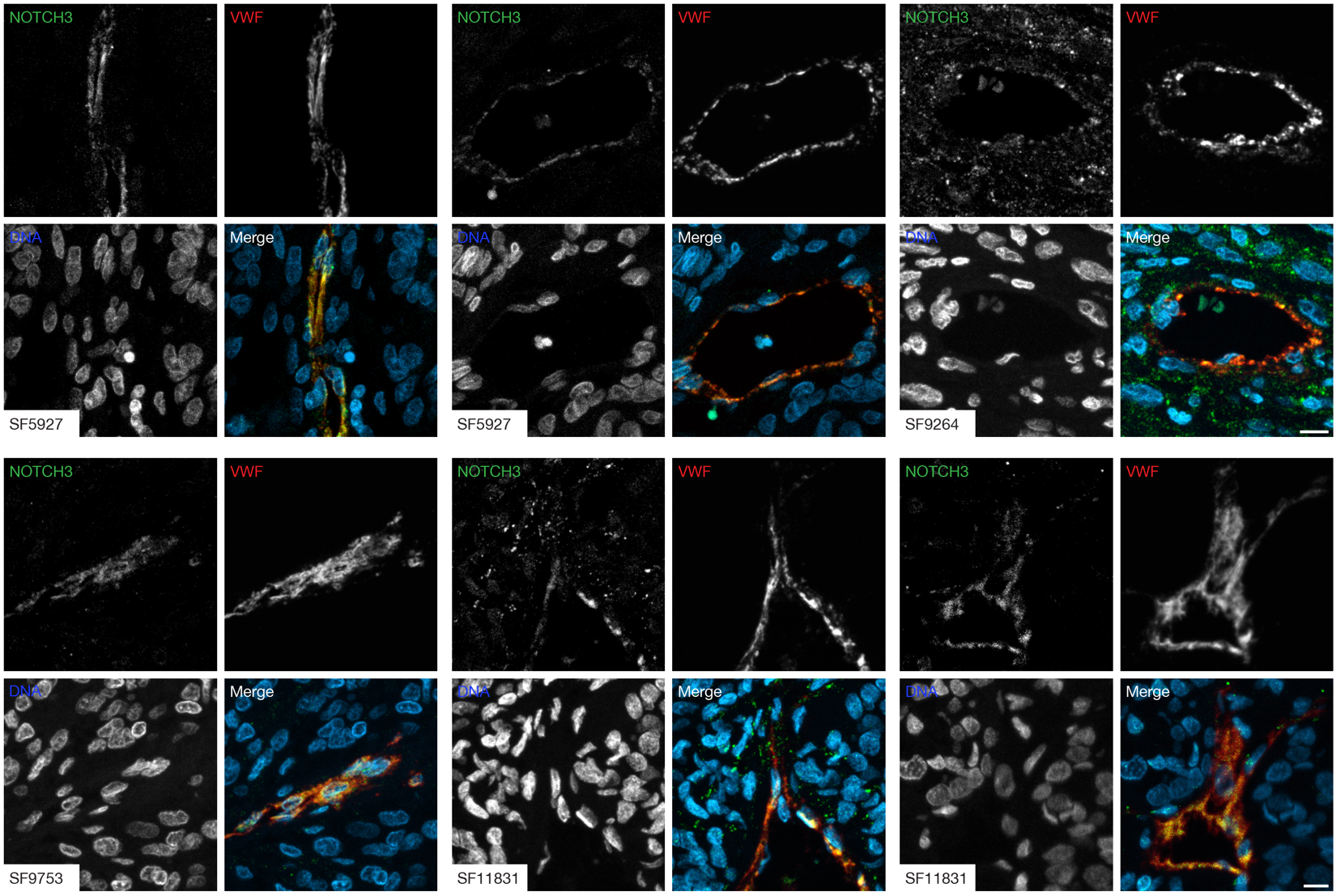
**Extended Data Fig. 9. WHO grade 3 meningioma immunofluorescence microscopy for NOTCH3 and the endothelial cell marker VWF.** NOTCH3 is expressed in and out of the perivascular niche, and is adjacent to endothelial cells in the perivascular niche, in meningiomas with WHO grade 3 histology. SF numbers indicate individual meningiomas. WHO grade defined using histological criteria. Scale bars, 10µm.

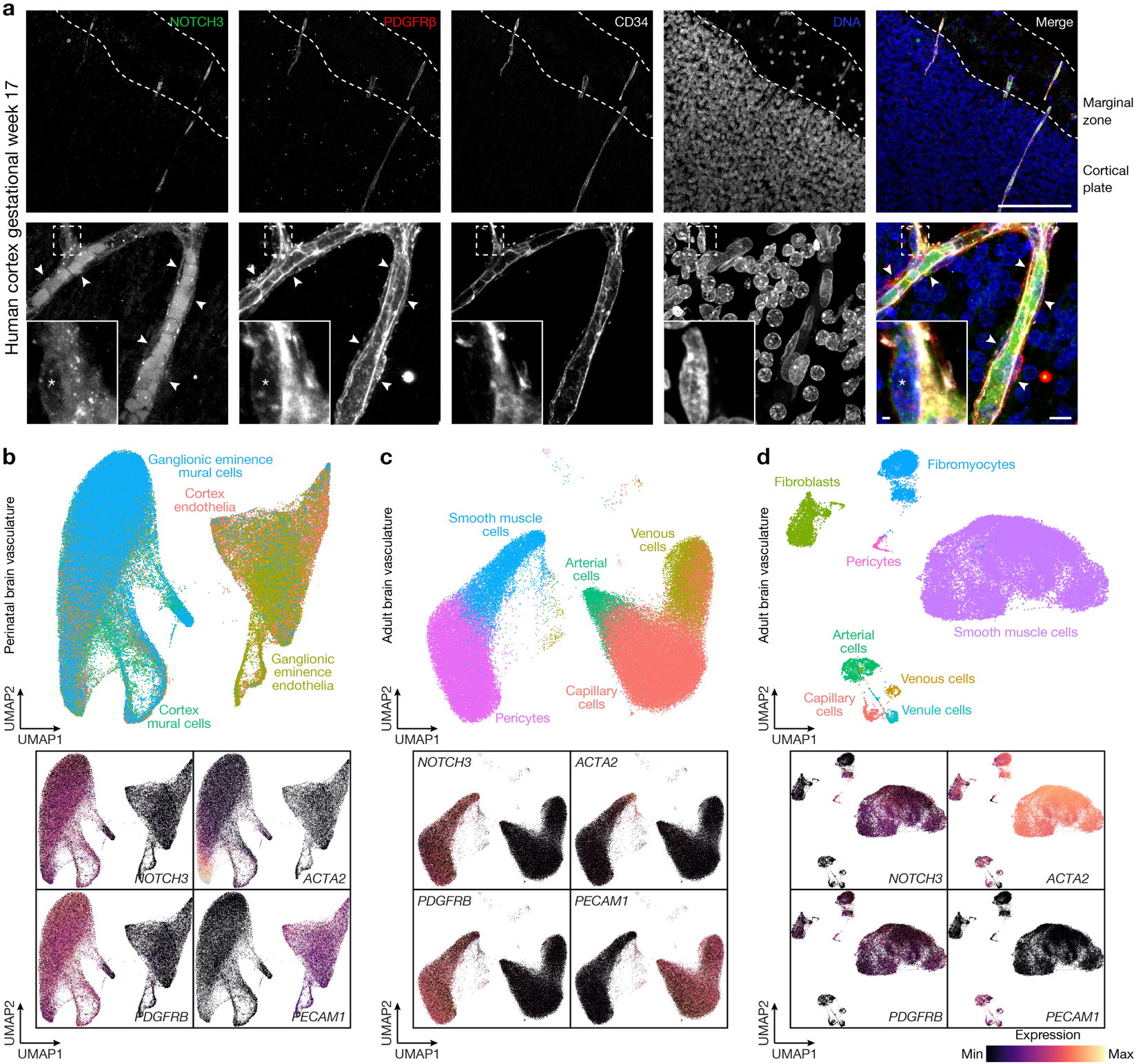
**Extended Data Fig. 10.** **NOTCH3 is expressed in mural cells during brain vascular development and homeostasis.** **a**, IF for NOTCH3, the mural cell marker PDGFRβ, and the endothelial cell marker CD34 in the developing human cortex from gestational week 17. NOTCH3 colocalizes with PDGFRβ in the marginal zone, which contributes to meningeal development^1–4^. Representative of n=3 biological replicates. DAPI marks DNA. Unperfused red blood cells in the vascular lumen are autofluorescent in the NOTCH3 channel. Scale bars, 100µm (top), 10µm (bottom), and 1µm (insert). **b**, Single-cell RNA sequencing cell cluster UMAP and feature plots from re-analysis of 139,134 perinatal human brain vasculature transcriptomes^5^ showing *NOTCH3* is enriched in developing mural cells marked by *ACTA2* or *PDGFRB* but not in developing endothelia marked by *PECAM1*. Samples for single-cell RNA sequencing were obtained from gestational weeks 15, 17, 18, 20, 22, and 23. **c**, Single-cell RNA sequencing cell cluster UMAP and feature plots from re-analysis of 84,138 adult human brain vasculature transcriptomes^6^ showing *NOTCH3* is expressed in mural cell lineages (pericytes, smooth muscle cells) but not endothelial cell lineages (venous cells, arterial cells, capillary cells). **d**, Single-cell RNA sequencing cell cluster UMAP and feature plots from re-analysis of 52,023 adult human brain vasculature transcriptomes^7^ showing *NOTCH3* is expressed in mural cell lineages (pericytes, smooth muscle cells, fibromyocytes) but not endothelial cell lineages (venous cells, arterial cells, capillary cells, venule cells).

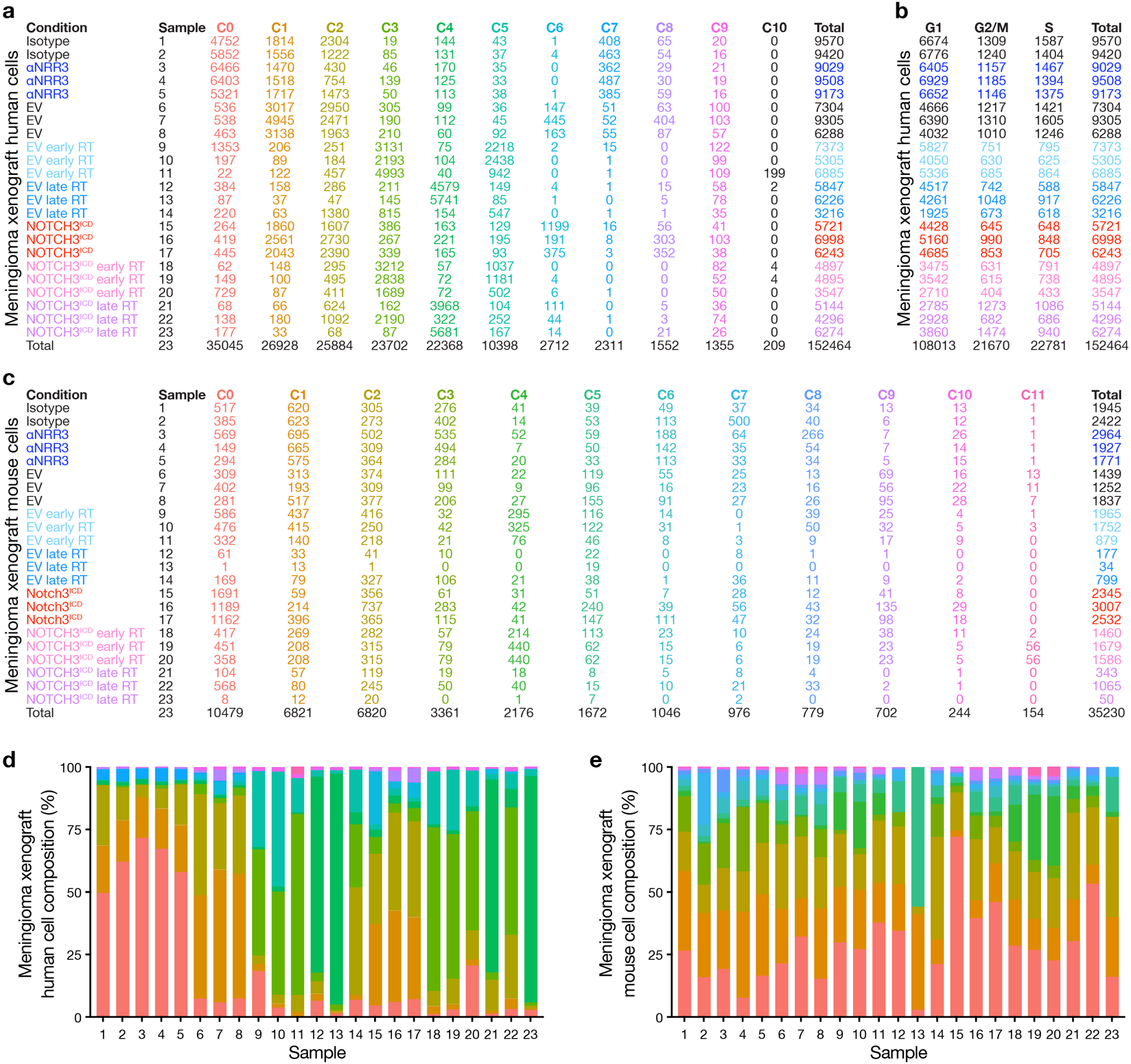
**Extended Data Fig. 11. Single-cell RNA sequencing of human and mouse cells from meningioma xenografts. a**, Single-cell counts across experimental conditions, biological replicates (samples), and UMAP clusters from CH-157MN meningioma xenograft human cells analyzed using single-cell RNA sequencing. **b**, Single-cell counts across experimental conditions, biological replicates, and phases of the cell cycle from meningioma xenograft human cells analyzed using single-cell RNA sequencing. Colors and rows (samples) as in **a**. **c**, Single-cell counts across experimental conditions, biological replicates, and UMAP clusters from meningioma xenograft mouse cells analyzed using single-cell RNA sequencing. **d**, Stacked bar plots showing the distribution of meningioma xenograft human single-cell RNA sequencing clusters across biological replicates. Colors as in **a**. **e**, Stacked bar plots showing the distribution of meningioma xenograft mouse single-cell RNA sequencing clusters across biological replicates. Colors as in **c**.

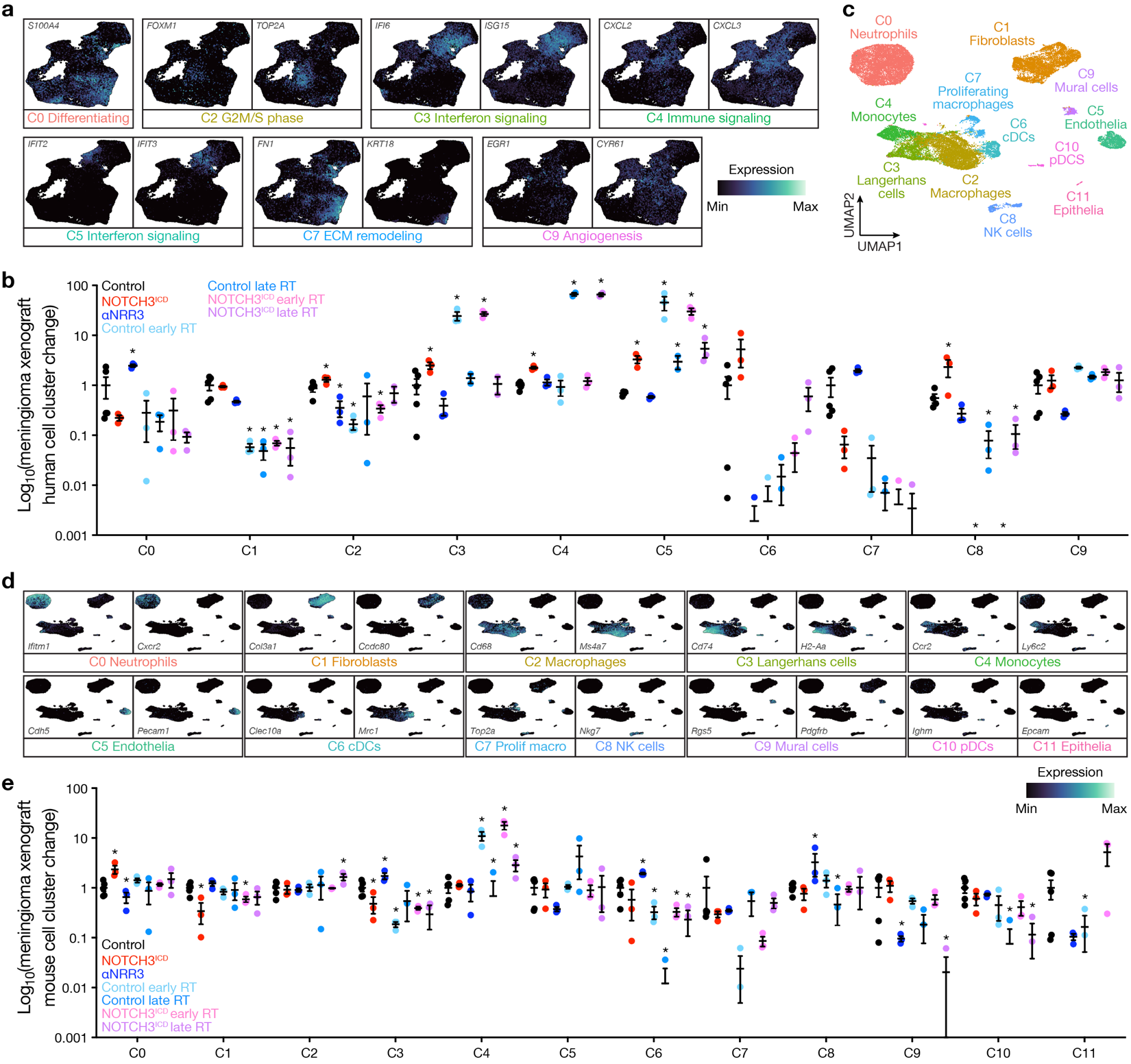
**Extended Data Fig. 12. Analysis of human and mouse cell types from single-cell RNA sequencing of meningioma xenografts. a**, Feature plots showing differentially expressed genes across UMAP clusters from CH-157MN meningioma xenograft human cells analyzed using single-cell RNA sequencing. See also Supplementary Table 5. **b**, Meningioma xenograft human single-cell cluster changes across experimental conditions and biological replicates. Student’s t tests. **c**, Single-cell RNA sequencing UMAP of 34,902 meningioma xenograft mouse cell transcriptomes showing microenvironment cell types. **d**, Feature plots showing differentially expressed genes across UMAP clusters from meningioma xenograft mouse cells analyzed using single-cell RNA sequencing. **e**, Meningioma xenograft mouse single-cell cluster changes across experimental conditions and biological replicates. See also Supplementary Table 6. Student’s t tests. *p<0.05.

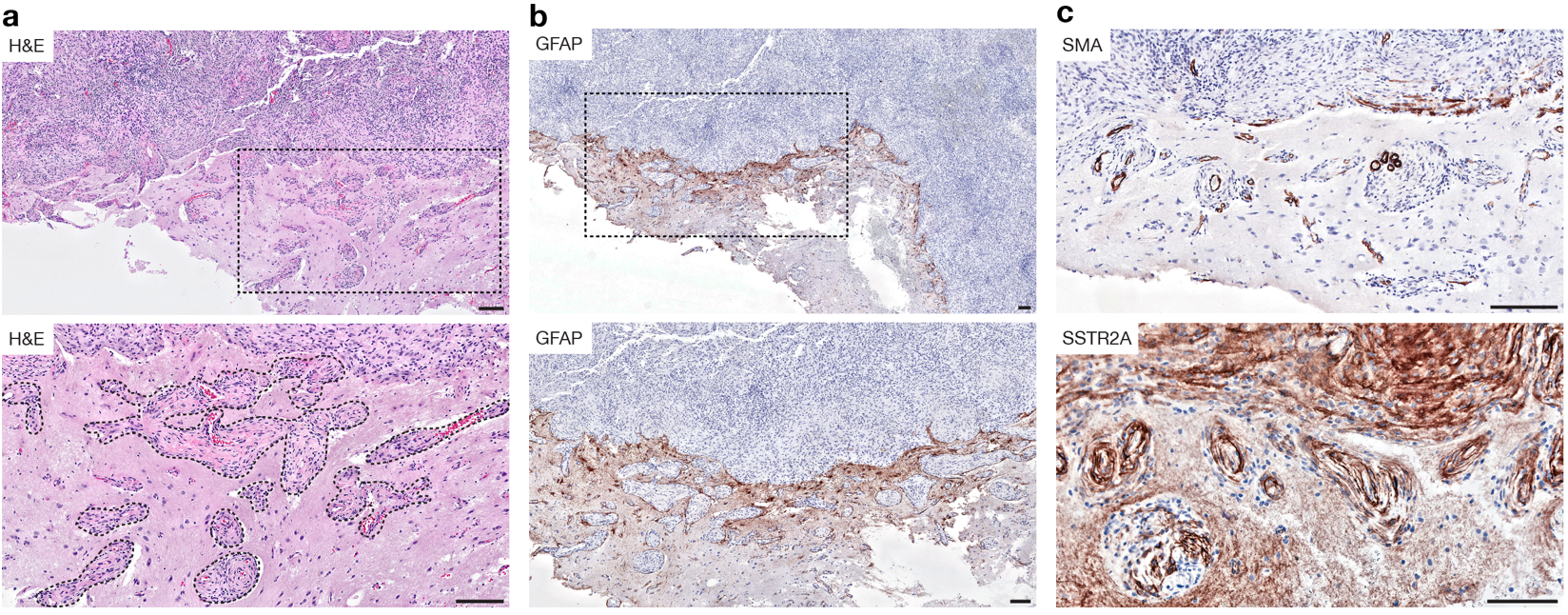
**Extended Data Fig. 13. Meningioma invasion of Virchow-Robin spaces. a**, H&E low magnification (top) and high magnification (box, bottom) images of human meningioma invasion into perivascular fluid-filled cavities surrounding perforating vasculature of the brain. Densely cellular meningioma is shown at the top of each image, and islands of perivascular tumor within the brain parenchyma (dashed lines, bottom) are shown at the bottom of each image. Scale bars, 100µm. **b**, Low magnification (top) and high magnification (box, bottom) images of IHC for the brain parenchyma marker GFAP validates invasion of unlabeled meningioma cells into Virchow-Robin spaces without direct invasion of the brain parenchyma itself. **c**, IHC for the mural cell marker SMA (top) or the meningioma cell marker SSTR2A (bottom) validates meningioma invasion into Virchow-Robin spaces.

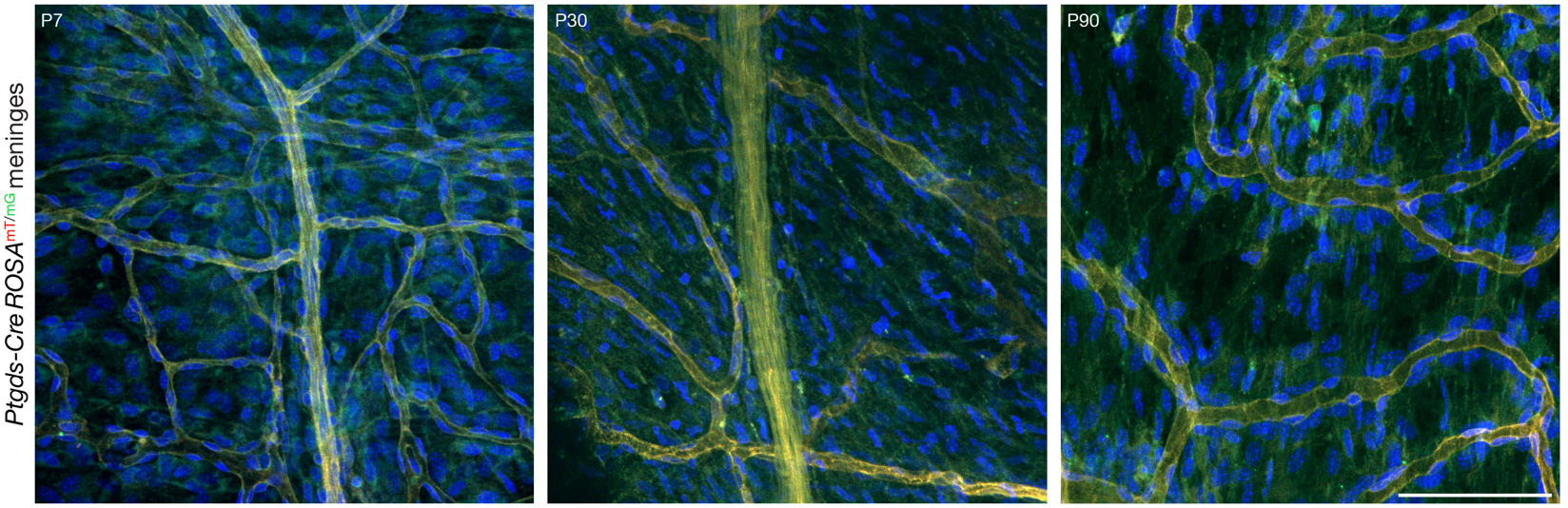
**Extended Data Fig. 14. *Ptgds* is expressed throughout the meninges.** Confocal microscopy of whole mount mouse convexity meningeal samples at P7, P30, or P90 after recombination of the *ROSA*^mT/mG^ allele using *Ptgds-Cre* shows PTGDS cells are diffusely expressed throughout perivascular and non-perivascular meningeal cells. DAPI marks DNA. Representative of n=3 biological replicates per timepoint. Scale bar, 100µm.

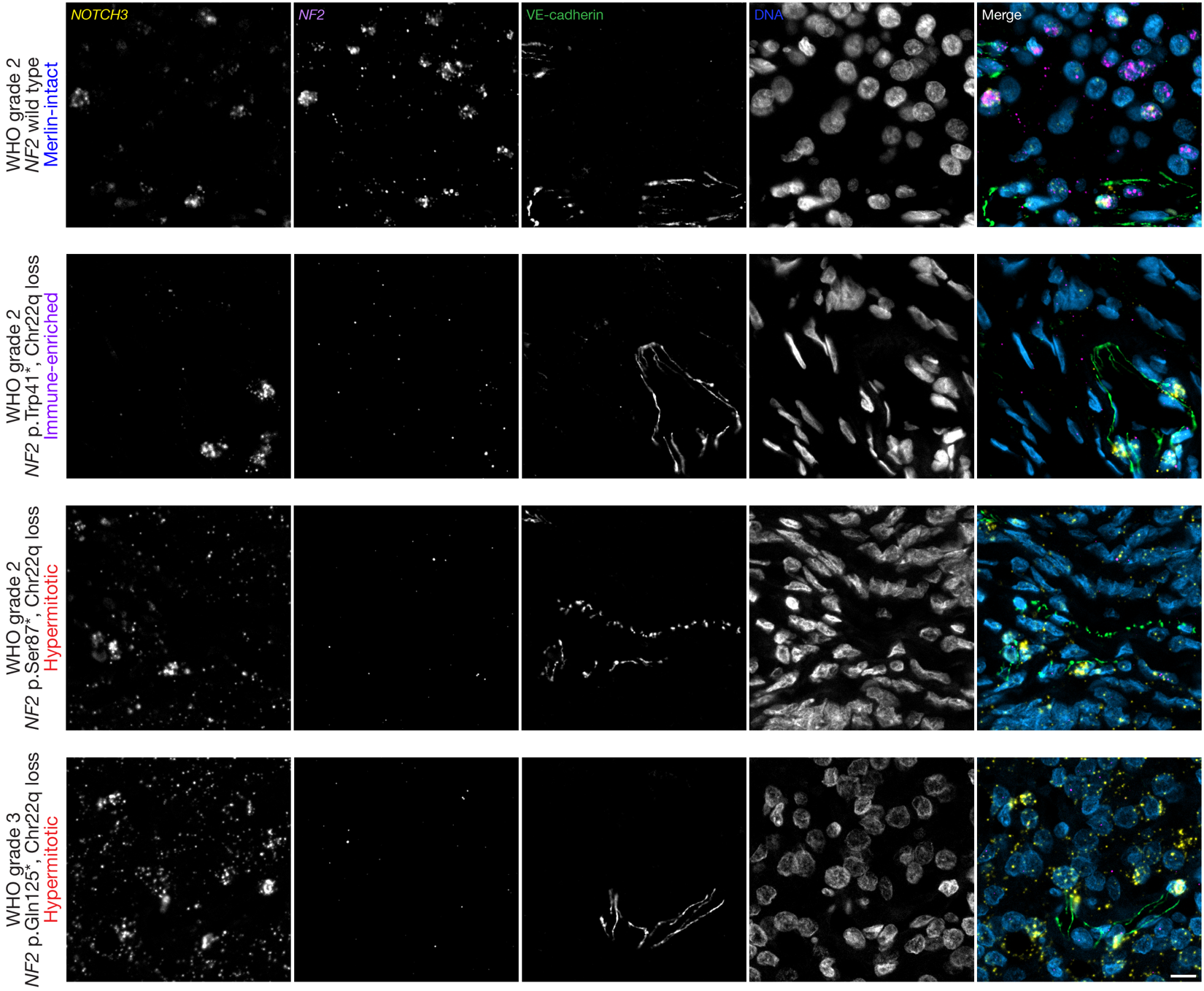
**Extended Data Fig. 15. *NOTCH3* RNAScope across meningioma DNA methylation groups.** RNAScope for *NOTCH3* and *NF2* integrated with IF for the endothelial cell marker VE-cadherin in meningiomas with paired DNA methylation profiling and targeted next-generation DNA sequencing reveals *NOTCH3* expression is enriched in meningiomas with biallelic *NF2* inactivation from DNA methylation groups with adverse clinical outcomes. WHO grade defined using histological criteria. Scale bar, 10µm.

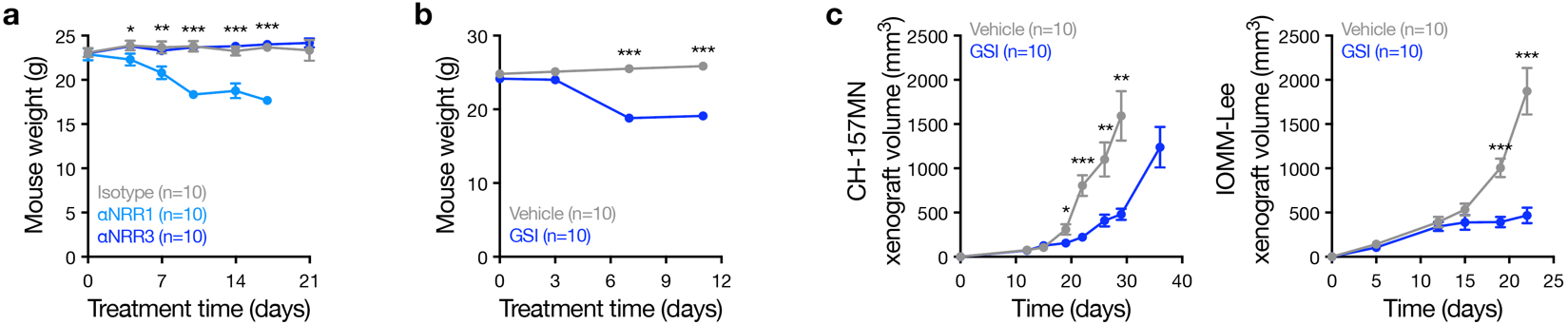
**Extended Data Fig. 16. NOTCH3 inhibition blocks meningioma xenograft growth without causing toxicity. a**, A NOTCH1 negative regulatory region neutralizing antibody (αNRR1), but not αNRR3, causes diarrhea and rash resulting in weight loss in mice harboring meningioma xenografts. Antibodies were delivered using biweekly IP injection. Student’s t tests. **b**, The γ-secretase inhibitor LY-411575 (GSI) delivered using daily IP injection causes diarrhea and rash resulting in weight loss in mice harboring meningioma xenografts. Student’s t tests. **c**, LY-411575 attenuates the growth of CH-175MN (left) or IOMM-Lee (right) meningioma xenografts in mice. Student’s t tests. Lines represent means and error bars represent standard error of means. *p<0.05, **p≤0.01, ***p≤0.0001.

**Extended references**
